# The α1 subunit of the Na,K-ATPase acts upstream of PI(4,5)P_2_ facilitating unconventional secretion of Fibroblast Growth Factor 2 from tumor cells

**DOI:** 10.1101/827691

**Authors:** Cyril Legrand, Roberto Saleppico, Jana Sticht, Fabio Lolicato, Hans-Michael Müller, Sabine Wegehingel, Eleni Dimou, Julia P. Steringer, Helge Ewers, Ilpo Vattulainen, Christian Freund, Walter Nickel

## Abstract

Fibroblast Growth Factor 2 (FGF2) is a tumor cell survival factor that is exported from cells by an unconventional secretory pathway. This process is based on direct translocation of FGF2 across the plasma membrane. FGF2 membrane translocation depends on PI(4,5)P_2_-induced formation of membrane-inserted FGF2 oligomers followed by extracellular trapping of FGF2 at the outer leaflet mediated by cell surface heparan sulfate proteoglycans. Beyond the well-characterized core mechanism of FGF2 membrane translocation, the Na,K-ATPase has been proposed to play a so far unknown role in unconventional secretion of FGF2. Here, we define a direct physical interaction of FGF2 with a subdomain of the cytoplasmic part of the α1 subunit of the Na,K-ATPase. Employing NMR spectroscopy and molecular dynamics simulations, we identified two lysine residues on the molecular surface of FGF2 that are shown to be essential for its interaction with α1. In intact cells, the corresponding lysine-to-glutamate variants of FGF2 were characterized by inefficient secretion and reduced recruitment to the inner plasma membrane leaflet as shown by single molecule TIRF microscopy. Our findings suggest that α1 acts upstream of PI(4,5)P_2_ facilitating efficient membrane translocation of FGF2 to the cell surface of tumor cells.

## Introduction

In eukaryotes, the majority of extracellular proteins is secreted through the ER/Golgi dependent secretory pathway ^1–4^. However, eukaryotic cells evolved additional mechanisms of protein transport into the extracellular space that have collectively been termed ‘unconventional protein secretion’ ^5, 6^. One of the most prominent examples for proteins secreted by unconventional means is Fibroblast Growth Factor 2 (FGF2) ^7^. FGF2 is a cell survival factor involved in tumor-induced angiogenesis with a broad significance for malignancies of both solid and hematological cancers ^8, 9^. Following secretion from tumor cells and their cellular microenvironment, FGF2 exerts its biological functions through both autocrine and paracrine signaling mediated by the formation of ternary complexes with FGF high affinity receptors and heparan sulfates on the surface of target cells.

Despite exerting its biological function in the extracellular space, FGF2 lacks a signal peptide and, therefore, does not have access to the classical, ER/Golgi dependent secretory pathway ^5, 6, 10, 11^. Based upon biochemical reconstitution experiments and bulk measurements of FGF2 secretion from cells, the unconventional secretory mechanism of FGF2 has been shown to be mediated by direct translocation of FGF2 across the plasma membrane ^12–14^. This process has recently been visualized in real time in living cells employing single molecule TIRF microscopy ^12^. Unconventional secretion of FGF2 depends on interactions with the α1 subunit of the Na,K-ATPase (α1) ^15^, Tec kinase ^16, 17^ and the phosphoinositide PI(4,5)P_2_ ^17–19^ at the inner plasma membrane leaflet as well as heparan sulfate chains of proteoglycans at the outer leaflet ^20, 21^. Consistently, residues in FGF2 that mediate interactions with PI(4,5)P_2_ [K127, R128 and K133; ^18, 22^] and heparan sulfates [K133; ^20, 22^] as well as the residue that is phosphorylated by Tec kinase [Y81; ^16, 17^] have been identified and shown to be critical for efficient secretion of FGF2. In addition, two cysteine residues (C77 and C95) on the molecular surface of FGF2 have been demonstrated to play a critical role in PI(4,5)P_2_ dependent formation of membrane inserted FGF2 oligomers ^23^. The latter have been shown to represent dynamic intermediates of FGF2 membrane translocation ^5, 7^ explaining the observation that FGF2 retains a fully folded state during membrane translocation to the cell surface ^24, 25^. Recently, key steps of the core mechanism of FGF2 membrane translocation have been reconstituted using an inside-out membrane model system based upon giant unilamellar vesicles along with entirely purified components ^22^. Based on the combined findings summarized above, a model of FGF2 membrane translocation was put forward with membrane-inserted FGF2 oligomers being key intermediates. These are assembled in a PI(4,5)P_2_ dependent manner at the inner leaflet and form membrane-spanning complexes. To complete FGF2 membrane translocation to the cell surface, FGF2 oligomers become disassembled at the outer plasma membrane leaflet mediated by membrane proximal heparan sulfates ^10, 11^. This model provides a compelling mechanism for directional transport of FGF2 from the cytoplasm into the extracellular space ^5, 7^. The general mechanism of this pathway of unconventional secretion is also relevant to other unconventionally secreted proteins ^5, 6^. For example, a role for PI(4,5)P_2_has been reported for unconventional secretion of Tau, HIV-Tat and Interleukin 1β ^26–31^. In addition, in case of Tau, sulfated glycosaminoglycans on cell surfaces have been shown to have a critical function in trans-cellular spreading that is mediated by unconventional secretion of Tau from donor cells ^27, 32^.

While the core mechanism of FGF2 membrane translocation is understood in great detail, the role of additional factors in unconventional secretion of FGF2 is unclear. In previous studies, a role for the catalytic α1-subunit of the plasma-membrane-resident sodium-potassium pump (Na,K-ATPase) in unconventional secretion of FGF2 has been proposed. Initial evidence was based on the observation that ouabain, an inhibitor of the Na,K-ATPase, blocks FGF2 transport into the extracellular space ^33–35^. These findings were corroborated by the observation that an α1 mutant to which ouabain cannot bind is capable of restoring FGF2 secretion in the presence of ouabain ^36^. Finally, α1 was one of the strongest hits in a genome-wide RNAi screen for gene products whose down-regulation inhibits FGF2 secretion from cells ^15^. Interestingly, α1 has also been suggested to be involved in unconventional secretion of HIV-Tat ^37^. However, as opposed to the detailed insight that is available about the core process of FGF2 membrane translocation, the precise function of α1 subunit of the Na,K-ATPase in unconventional secretion of FGF2 is unknown ^5, 7, 38, 39^.

In the current study, we define a sub-domain in the cytoplasmic part of α1 that forms the binding surface for FGF2. Employing a chemical crosslinking approach, we identified a physical 1:1 complex of FGF2 and this minimal α1 domain as the main crosslinking product. Using NMR spectroscopy, we identified two lysine residues (K54 and K60) on the molecular surface of FGF2 that are critical for binding to α1. Using both docking studies and molecular dynamics simulations, we validated the role of K54 and K60 in a thermodynamically relevant model system. Consistently, FGF2 mutants lacking K54/K60 were found to be largely incapable of binding to α1 as shown by two different kinds of biochemical protein-protein interaction assays. These observations were further analyzed in cell-based secretion assays, demonstrating a drop in FGF2 secretion efficiency when K54 and K60 were replaced by glutamates. Using a recently established single molecule TIRF microscopy assay designed to quantify FGF2 recruitment at the inner plasma membrane leaflet in living cells ^12^, we found that K54/60E mutants of FGF2 failed to get efficiently recruited at the inner leaflet of the plasma membrane. By contrast, a FGF2 variant form lacking the ability to bind to PI(4,5)P_2_ was found at the inner leaflet at amounts comparable to the wild-type protein. These results demonstrate that FGF2 binding to α1 precedes recruitment of FGF2 by PI(4,5)P_2_. Furthermore, as opposed to *in vitro* experiments demonstrating efficient binding of FGF2 to PI(4,5)P_2_ with a K_D_ of about 1 µM ^17–19, 22^, these findings suggest that, in the context of intact cells, α1 is required for efficient binding of FGF2 to PI(4,5)P_2_. We propose that α1 is an auxiliary factor that acts upstream of PI(4,5)P_2_ to increase the efficiency of unconventional secretion of FGF2 by accumulating FGF2 at the inner plasma membrane leaflet.

## Results

### Identification of a sub-domain in the cytoplasmic part of the α1 subunit of the Na,K-ATPase that directly interacts with FGF2

While we established an interaction of FGF2 with the large cytoplasmic domain of the α1 subunit of the Na,K-ATPase in a previous study ^15^, the starting point of the current investigation was to define a minimal sub-domain that contains the protein interaction surface for FGF2. In Fig. 1A, the set of GST fusion proteins used in this study is depicted carrying different forms of the cytoplasmic domain of α1. They contained either the complete (GST-α1-CD1-3) or various sub-domains of the cytoplasmic domain of α1 as indicated. Based upon a structural analysis, we identified a globular domain of about 20 kDa in α1 that contains a highly acidic molecular surface as a potential recruitment site for FGF2. This sub-domain is contained in the third loop of the cytoplasmic domain of α1 (α1-subCD3) that is almost identical to the N-domain of the α1 subunit of the Na,K-ATPase ^34, 40^. In a first set of experiments, a biochemical pull-down approach was used to quantify the interaction of FGF2 with the various GST-α1 fusion proteins (Fig. 1B). The construct containing the complete cytoplasmic domain of α1 [α1-CD1-3; ^15^] was used as a positive control. These experiments revealed that FGF2 binds with similar efficiency to α1-CD1-3, α1-CD3 (containing only the third loop of the cytoplasmic domain of α1) and the small sub-domain of loop 3 (α1-subCD3). By contrast, FGF2 did neither bind to a significant extent to a construct lacking the sub-domain of loop 3 (GST-α1-CD3-ΔsubCD3) nor to GST alone used as a negative control (Fig. 1C).

**Fig. 1:**
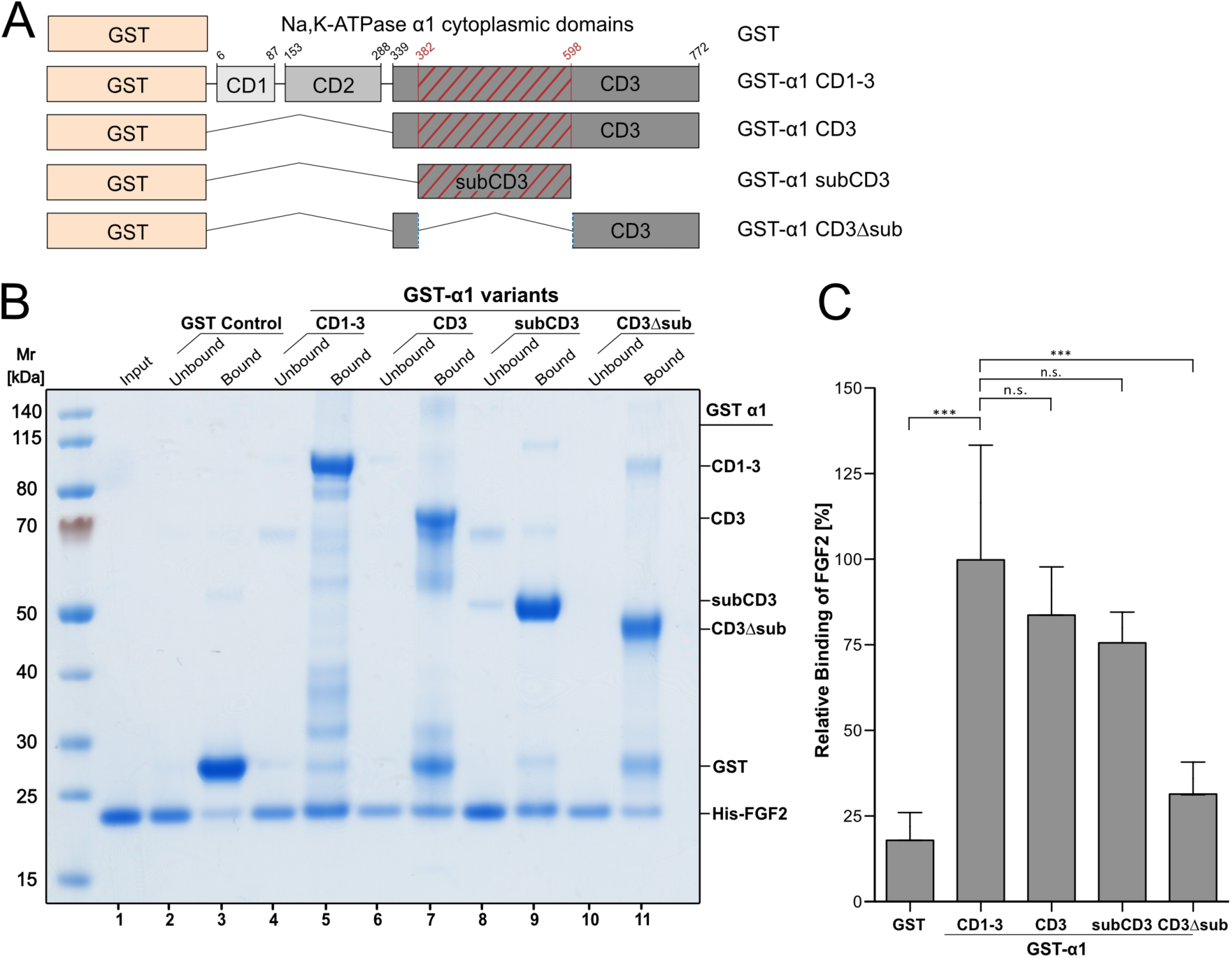
Identification of a sub-domain in the cytoplasmic domain of α1 that is both necessary and sufficient for binding of FGF2. A) Schematic representation of GST-tagged α1 constructs used in this study. B) Biochemical pull-down experiments of FGF2 using GST fusion proteins of various versions of the cytoplasmic domain of α1. Lane 1: FGF2 input (2.5%); Lanes 2-3: GST control; Lanes 4-5: GST-α1-CD1-3; Lanes 6-7: GST-α1-CD3: Lanes 8-9: GST-α1-subCD3; Lanes 10-11: GST-α1-CD3Δsub. Bound (33% of each fraction) and unbound material (2.5% of each fraction) was analyzed by SDS-PAGE and Coomassie Blue protein staining. The results shown are representative for five independent experiments. C) The intensity of the FGF2 protein bands from SDS PAGE (panel B) was analyzed with the ImageStudio software package (LI-COR Biosciences). Ratios of bound versus unbound FGF2 were calculated and normalized to the amounts of FGF2 bound to GST-α1-CD1-3 containing the complete cytoplasmic domain of α1. Data were analyzed from five biological replicates and a one-way analysis of variance was performed (n.s. = p-value > 0.05; *** = p-value ≤ 0.001).

The results from biochemical pull-down experiments were confirmed employing an AlphaScreen^®^ protein-protein interaction assay (Fig. 2). Cross-titration experiments using a wide range of combinations of concentrations of FGF2 and various α1 constructs demonstrated that FGF2 binds to the complete cytoplasmic domain (α1-CD1-3), loop 3 alone (α1-CD3) and α1-subCD3, the sub-domain of loop 3 of the cytoplasmic domain (Fig. 2A and quantified in Fig. 2B). By contrast, FGF2 did not bind to either α1-CD3-ΔsubCD3 or GST alone (Fig. 2A and 2B). Using an untagged version of FGF2 as a competitor for the interaction of His-tagged FGF2 and GST-tagged α1 constructs, the AlphaScreen^®^ assay also allowed for a quantitative comparison of the affinities between FGF2 and α1-CD1-3, α1-CD3 as well as α1-subCD3, respectively [Fig. 2C; ^15, 41^]. As reported previously, FGF2 binds to the complete cytoplasmic domain of α1 with an apparent K_D_ of about 0.8 µM ^15^. With 1.1, 1.3 and 0.8 µM, similar apparent K_D_s were found for α1-CD1-3, α1-CD3 and α1-subCD3, respectively (Fig. 2C), suggesting that the sub-domain of loop 3 in the cytoplasmic domain of α1 contains the full binding site that is both necessary and sufficient for FGF2 recruitment.

**Fig. 2:**
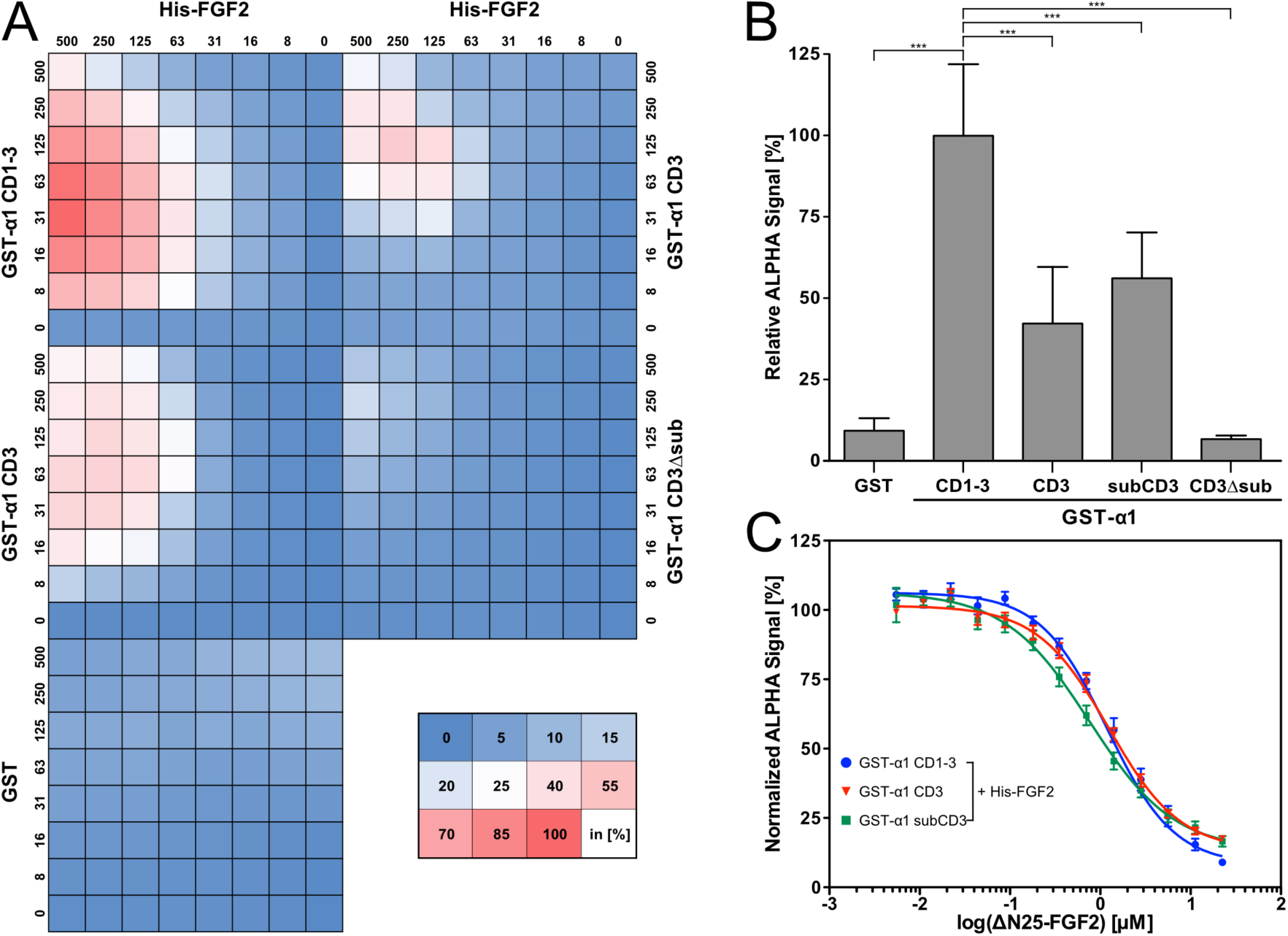
FGF2 binds to α1-subCD3 with sub-micromolar affinity as analyzed by an AlphaScreen protein-protein interaction assay. A) Cross-titration experiments conducted with His-tagged FGF2 and GST-tagged variant forms of the cytoplasmic domain of α1 as indicated. Data were normalized to the signal intensity measured at 250 nM His-tagged FGF2 and 63 nM GST-α1-CD1-3. The shown heat map is an average from five biological replicates. See Material and Methods for details. B) Quantification and statistical analysis of the relative Alpha signal intensity from the cross-titration experiment with His-tagged FGF2 and the GST-fusion proteins of the cytoplasmic domain of α1 as indicated in panel A. A one-way analysis of variance was performed to test for statistical significance of the observed differences (*** = p-value ≤ 0.001) C) Determination of affinity using IC50 values from competition assays. GST-tagged α1 variant forms (CD1-3, CD3 or subCD3) and His-tagged FGF2 proteins were mixed and subjected to a serial dilution of ΔN25-FGF2 used as an untagged competitor. Data analysis was done from six independent biological replicates.

To determine the stoichiometry of the interaction between the minimal FGF2 binding domain in the cytoplasmic part of α1 (α1-subCD3) and FGF2, we conducted cross-linking experiments (Fig. 3). As explained in Materials and Methods, two types of chemical crosslinkers were chosen targeting either cysteine (BMOE; Fig. 3A and 3C) or lysine (DSG; Fig. 3B and 3D) side chains. The formation of cross-linking products was monitored by both a Western analysis using anti-FGF2 and anti-α1 antibodies (Fig. 3A and 3B) and an SDS-PAGE analysis based upon Coomassie protein staining (Fig. 3C and 3D). For both read-outs and crosslinkers, the main product was characterized by a migration behavior that is consistent with a molecular weight of about 46 kDa. This product was recognized by both anti-FGF2 and anti-α1 antibodies (Fig. 3A and 3B), suggesting a 1:1 complex of FGF2 and the α1-subCD3. For BMOE in particular, substantial amounts of the 1:1 cross-linking product were formed as it was readily detectable by Coomassie protein staining as well (Fig. 3C). Additional crosslinking products with higher molecular weights were identified as well which form because of the ability of both FGF2 and α1 to form homo-oligomers (Fig. 3A-D; lanes 5 and 6). However, the formation of FGF2/α1 heterodimers occurs preferentially as they are formed at the expense of FGF2 homodimers (Fig. 3A-D, lane 7 versus lane 9). These results are consistent with the experiments shown in Figs. 1 and 2, establishing a direct physical contact between FGF2 and α1 with a heterodimer being the basic unit of this interaction.

**Fig. 3:**
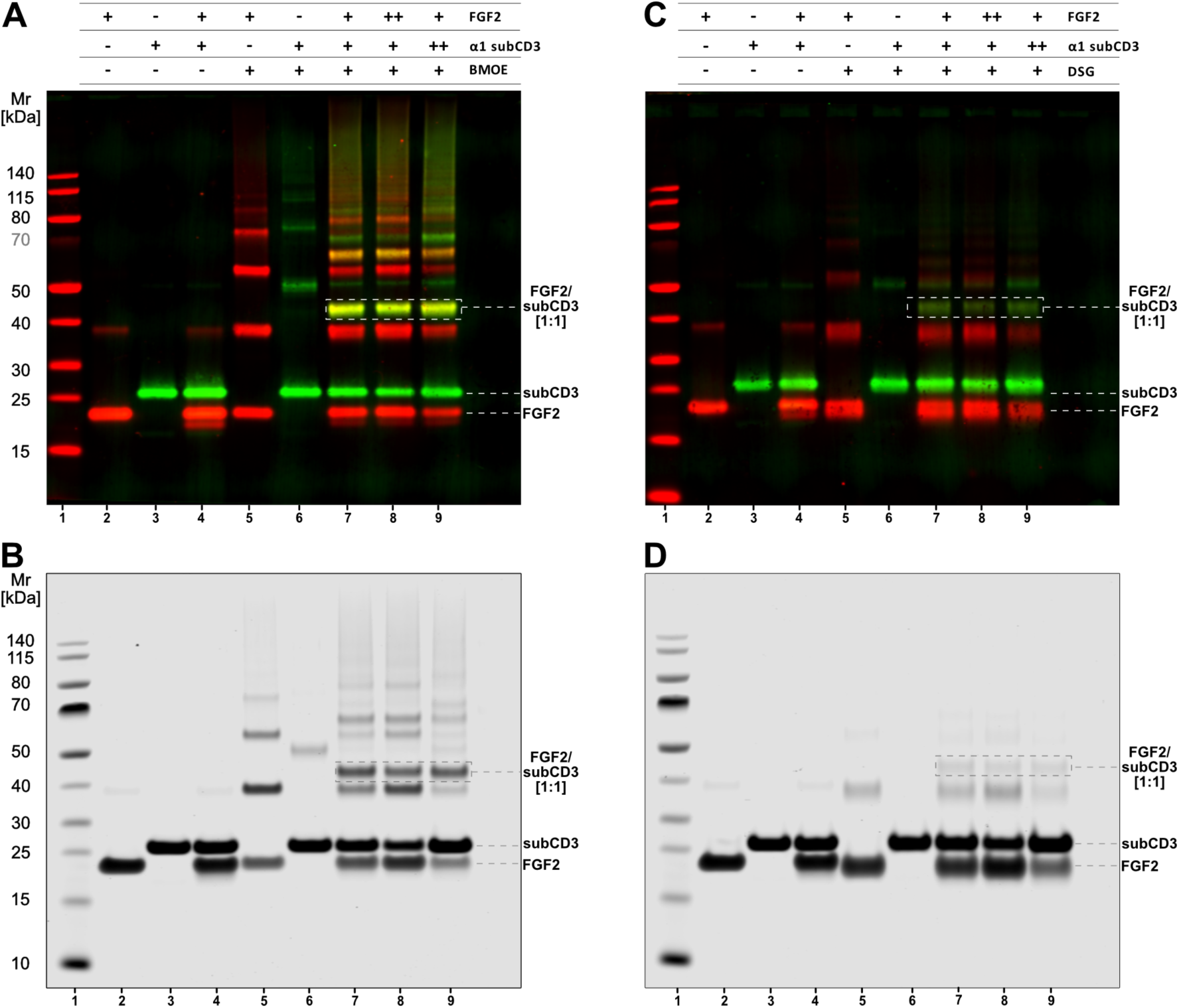
The basic binding unit between FGF2 and α1-subCD3 is a heterodimer as analyzed by chemical crosslinking experiments. FGF2 and α1-subCD3 were mixed at a total concentration of 20 µM at molar ratios of 1:1 (+/+), 1:2 (+/++) and 2:1 (++/+) (FGF2/α1-subCD3). As chemical crosslinkers, Bismaleidomethane (BMOE) and Disuccinimidyl glutarate (DSG) were used at the molar ratios provided in Materials and Methods. Samples were analyzed by SDS-PAGE followed by either Western blotting (panels A and B) or total protein staining using Coomassie. FGF2 and α1-subCD3 migrate with an apparent molecular weight of 18 kDa and 27 kDa, respectively. The crosslinking product with an apparent stoichiometry of 1:1 shows a migration behavior that corresponds to a molecular weight of about 45 kDa as indicated. Lane 1: Molecular weight standards; Lane 2: Input FGF2; Lane 3: Input α1-subCD3; Lane 4: FGF2 plus α1-subCD3 without crosslinker; Lane 5: FGF2 plus crosslinker; Lane 6: α1-subCD3 plus crosslinker; Lanes 7-9: FGF2 plus α1-subCD3 at 1:1, 2:1 and 1:2 stoichiometries in the presence of crosslinker. A) BMOE; Immunoblots using anti-FGF2 and anti-α1 primary antibodies B) DSG; Immunoblots using anti-FGF2 and anti-α1 primary antibodies C) BMOE; Total protein visualized by Coomassie staining D) DSG; Total protein visualized by Coomassie staining

### Identification of the FGF2 residues K54 and K60 as part of the binding interface with the α1 subunit of the Na,K-ATPase employing NMR spectroscopy

In order to map the binding interface of α1-subCD3 on FGF2, NMR experiments were performed using a FGF2 variant form (FGF2-C77/95S) that is incapable of oligomerization ^22, 23^. ^1^H-^15^N-HSQC spectra of ^15^N-labeled FGF2-C77/95S in the absence and in the presence of α1-subCD3 (in molar ratios of 1:1 and 1:2) were acquired and assignments were transferred from published data ^42^. Chemical shift differences upon addition of α1-subCD3 were negligible for assigned peaks (<0.04 ppm, Fig. 4-supplement 1) but some signals showed a clear reduction in peak intensity (Fig. 4 and Fig. 4-supplement 1). In agreement with the observed K_D_ value of about 0.8 µM, this speaks for a slow to intermediate exchange regime on the NMR timescale. The larger size of the complex compared to isolated FGF2 also manifests in a general line-broadening of signals that is seen as a small but significant reduction in peak intensities (Fig. 4-supplement 1). Three of the six residues with strong reduction in signal intensities localize to the region spanning K54 to K60. The two effected lysine residues K54 and K60 are exposed on the molecular surface of FGF2 and, therefore, were considered to potentially form a charged surface for the interaction with α1-subCD3.

**Fig. 4:**
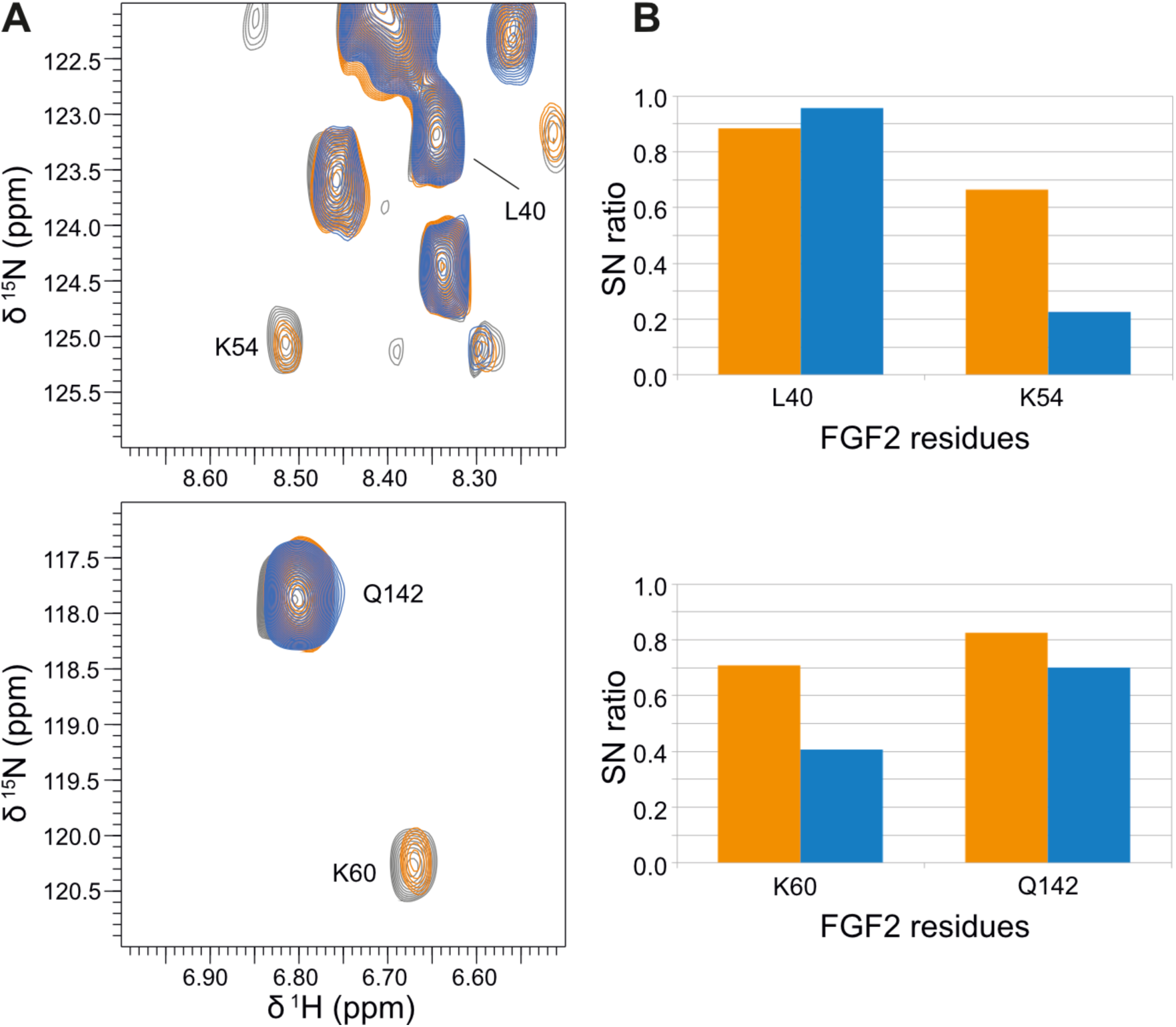
NMR analysis of the interaction between FGF2-C77/95S and α1-subCD3. A) Shown are two regions of the overlay of ^1^H-^15^N-HSQC spectra of 77 µM ^15^N-labeled FGF2-C77/95S in the absence (grey) or presence of 70µM (orange) or 138 µM ATP1A1-subCD3 (blue). B) Signal-to-noise ratios (SN ratio) for indicated peaks of the spectra shown in A are plotted here with SN(+70µM α1-subCD3)/SN(-α1-subCD3) in orange and SN(+138µM α1-subCD3)/SN(-α1-subCD3) in blue.

### Bioinformatical analysis of FGF2 versus FGF proteins with signal peptides

A principal approach to identify residues in FGF2 with a specific role in its unconventional secretory pathway is to compare the primary sequence of FGF2 with other FGF family members carrying signal peptides for ER/Golgi dependent protein secretion. In this way, in previous studies, we have identified two cysteine residues on the molecular surface of FGF2 (C77 and C95) that are absent from all FGF family members with signal peptides. In subsequent studies, these residues were demonstrated to play a critical role in unconventional secretion of FGF2 from cells ^12, 22, 23^. As illustrated in Fig. 5, similar to C77 and C95 (highlighted in yellow), the lysine residues in position 54 and 60 (highlighted in green) of FGF2 are absent from most FGF family members traveling through the ER/Golgi-dependent secretory pathway. By contrast, the majority of residues such as C33 and C100 are conserved throughout the FGF family irrespective of the mode of secretion of the respective FGF proteins, suggesting that they are important for the overall fold of FGF proteins. The identification of K54 and K60 by NMR spectroscopy as residues that potentially localize to the interface mediating FGF2 binding to the cytoplasmic domain of α1 is consistent with their absence from FGF family members that are secreted by the signal-peptide-dependent, classical secretory pathway and suggests a specific role in unconventional secretion of FGF2.

**Fig. 5:**
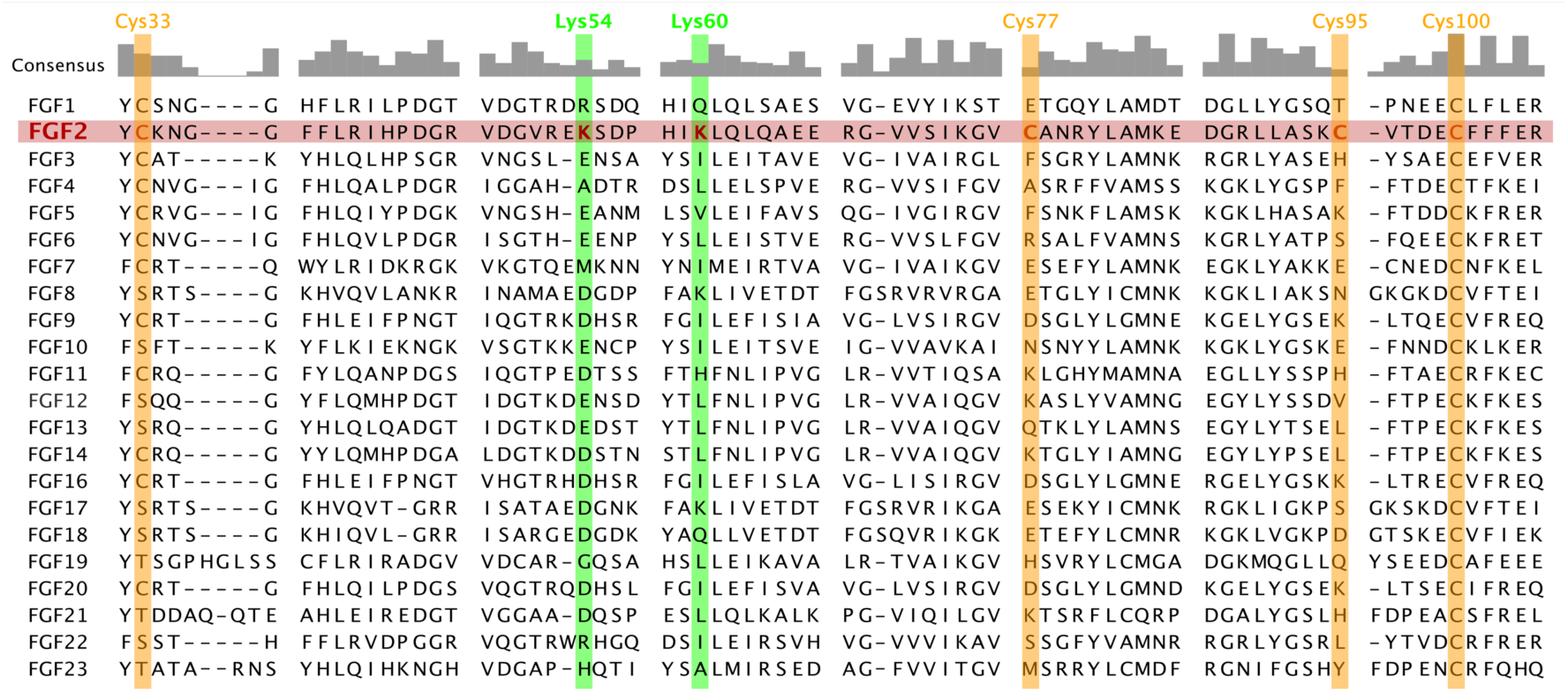
Residues critical for unconventional secretion of FGF2 are absent from most FGF family members carrying signal peptides for ER/Golgi-dependent protein secretion. A sequence alignment of the FGF protein family covering residues 32 to 105 from FGF2 was used to identify residues with a potential role in unconventional secretion of FGF2. Residues highlighted in yellow represent cysteine residues in FGF2 that are either fully conserved throughout the FGF family (C33 and C100) or are uniquely present in FGF2 (C77 and C95). Similar to C77 and C95, K54 and K60 highlighted in green are absent from most FGF family proteins carrying signal peptides.

### FGF2 mutant forms lacking K54 and K60 are impaired in binding to the cytoplasmic domain of the α1 subunit of the Na,K-ATPase

To test whether K54 and K60 in FGF2 are residues that are indeed critical for binding to the cytoplasmic domain of α1, we generated three FGF2 mutants in which these lysines were substituted by glutamate residues either separately or combined. In a first series of experiments, these FGF2 mutants were tested for binding to a GST fusion protein containing the α1-subCD3 domain employing biochemical pull-down experiments (Fig. 6). A representative example is given in Fig. 6A with the corresponding quantification and statistical analysis of four independent experiments shown in Fig. 6B. While both the FGF2-K54E and the FGF2-K60E mutants already showed reduced binding efficiency, the interaction of FGF2-K54/60E to α1-subCD3 was strongly impaired.

**Fig. 6:**
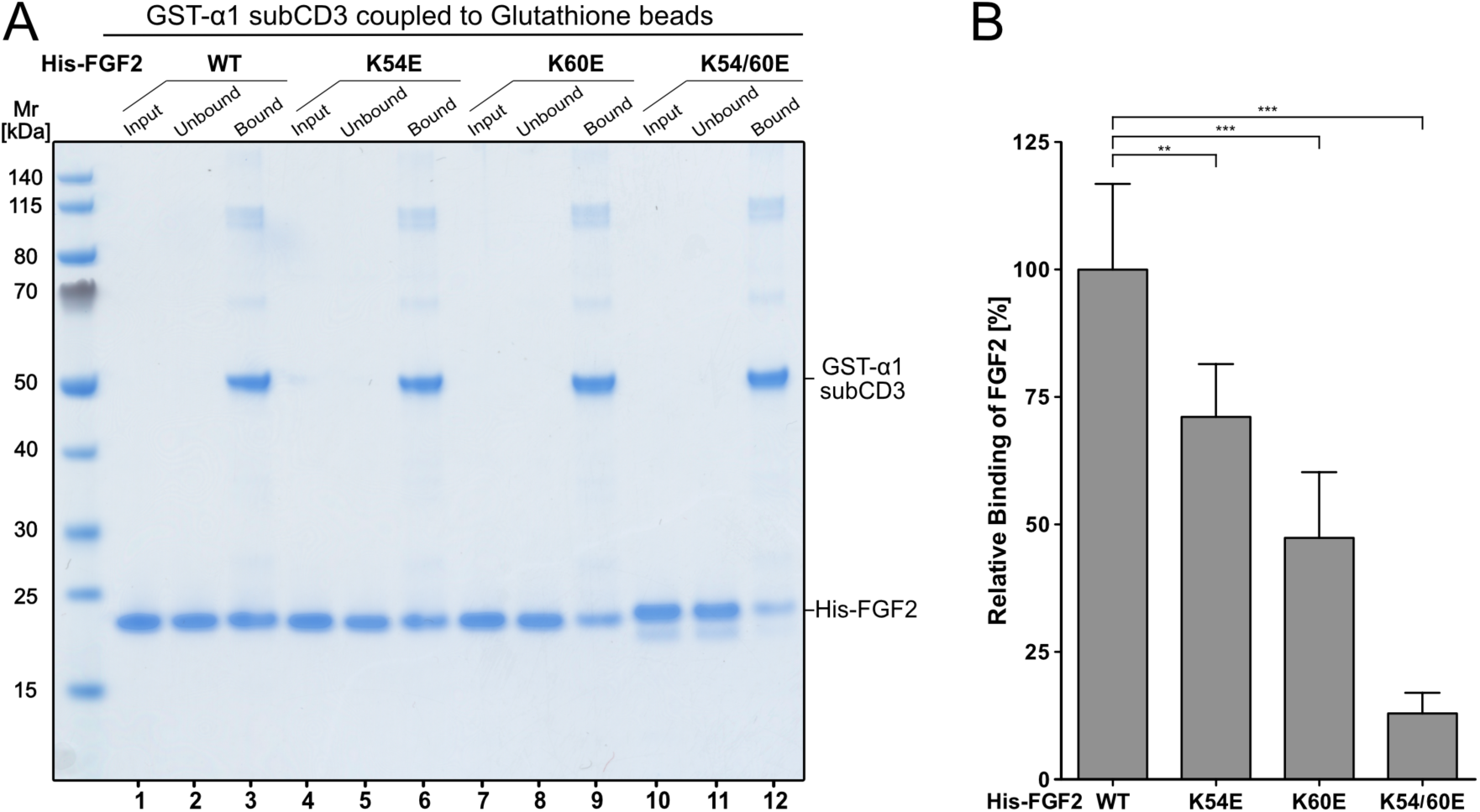
K54 and K60 are required for efficient binding of FGF2 to α1-subCD3 as shown by biochemical pull-down experiments. A) Glutathione beads containing GST-tagged α1-subCD3 were incubated with His-tagged forms of FGF2 as indicated. FGF2 input (2.5%), unbound material (2.5%) and bound FGF2 (33%) were analyzed by SDS-PAGE and Coomassie Blue protein staining. Lanes 1-3: FGF2-wt; Lanes 4-6: FGF2-K54E; Lanes 7-9: FGF2-K60E; Lanes 10-12: FGF2-K54/60E. For details see Material and Methods. B) Quantification and statistical analysis of relative binding efficiencies of FGF2-wt and mutant forms to α1-subCD3. The intensities of the FGF2 protein bands from panel A were analyzed with the ImageStudio software package (LI-COR Biosciences). Ratios of bound and unbound material were calculated and normalized to FGF2-wt. Data were analyzed from five biological replicates and a one-way analysis of variance was performed (n.s. = p-value > 0.05; ** = p-value ≤ 0.01; *** = p-value ≤ 0.001).

These results were confirmed using the AlphaScreen^®^ protein-protein interaction assay (Fig. 7). In panel A, a cross-titration experiment is shown analyzing a wide range of concentrations quantifying the interaction between α1-subCD3 and FGF2-wt, FGF2-K54E, FGF2-K60E as well as FGF2-K54/60E, respectively. Based upon the combination of concentrations resulting in the strongest interaction between FGF2-wt and α1-subCD3, four independent experiments were statistically evaluated as shown in Fig. 7B. Consistent with the biochemical pull-down experiments (Fig. 6), single amino acid substitutions of either K54 or K60 caused partial inhibition of FGF2 binding to α1-subCD3. Substitution of both lysines by glutamates (FGF2-K54/60E) resulted in an almost complete lack of binding (Fig. 7B).

**Fig. 7:**
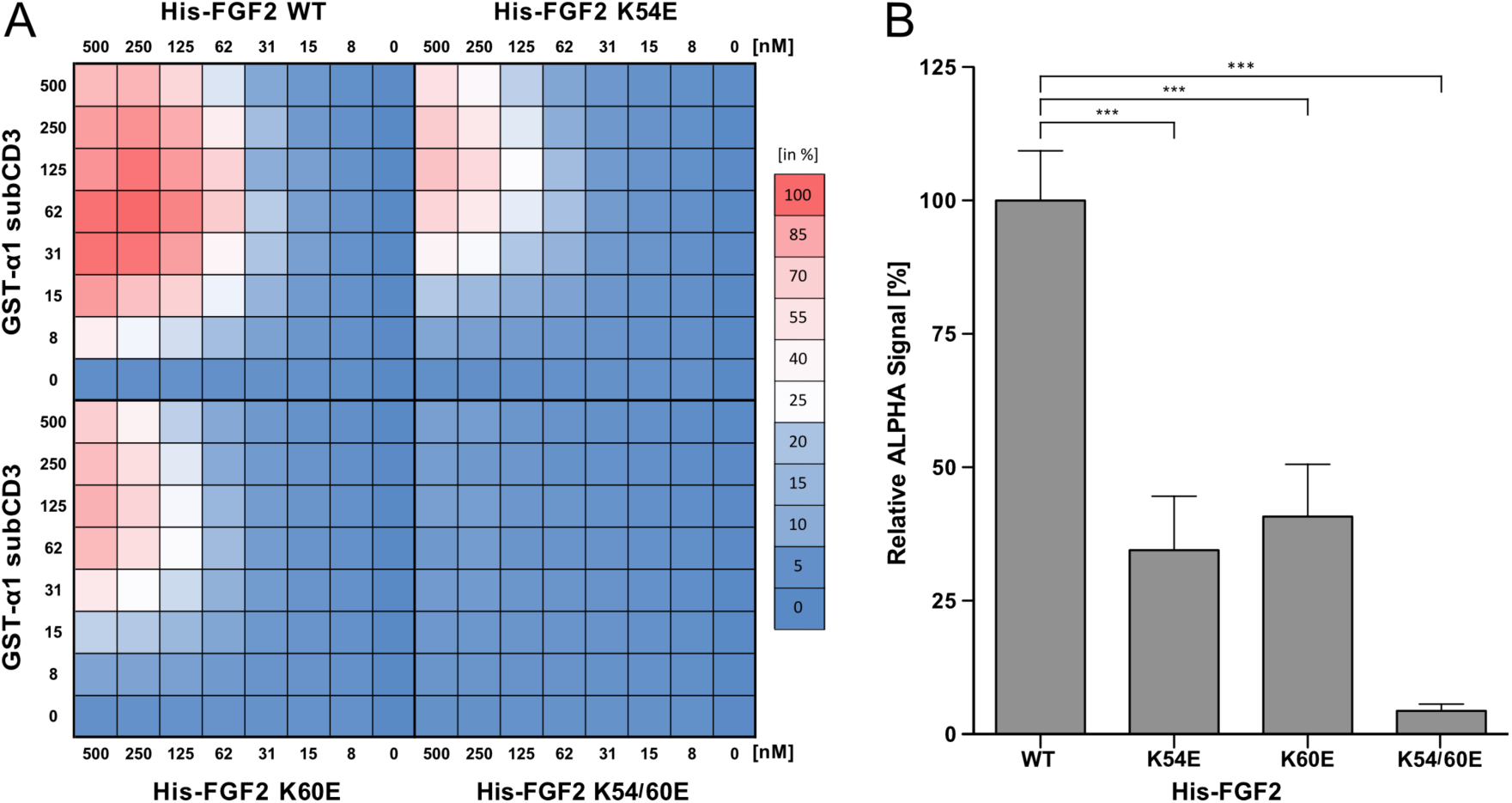
K54 and K60 are required for efficient binding of FGF2 to α1-subCD3 as analyzed by AlphaScreen protein-protein interaction experiments. A) Cross-titration experiments conducted with various forms of His-tagged FGF2 and GST-tagged α1-subCD3. For each biological replicate, data were normalized to the signal measured for His-FGF2/GST-α1-CD1-3 at 250nM and 63nM respectively. As indicated by the color legend, data were represented as a heat map with the highest signal set to 100% (displayed in red) and the lowest signal set to 0% (displayed in blue). The shown heat map is an average from five biological replicates. For details, see Material and Methods. B) Quantification and statistical analysis of relative Alpha signal intensities from the cross-titration experiment shown in panel A. Statistically significance was tested using a one-way analysis of variance (*** = p-value ≤ 0.001)

To test whether substitutions of K54 and K60 in FGF2 cause pleiotropic effects affecting other functional properties beyond binding to α1-subCD3, we compared all four variant forms of FGF2 (wt, K54E, K60E and K54/60E) with regard to their ability to bind to PI(4,5)P_2_ and to oligomerize (Fig. 7 - supplement 1, panel A), to bind to heparin (Fig. 7- supplement 1, panel B) as well as tested for potential folding defects based upon thermal stability (Fig. 7 - supplement 1, panel C). These experiments revealed that the substitution of K54 and K60 by glutamates does not have any impact on the parameters listed above. In conclusion, using two independent assays to quantify the interaction between FGF2 and α1-subCD3, the FGF2 residues K54 and K60 could be established as critical residues for the interaction of FGF2 with the cytoplasmic domain of α1.

### Analysis of the FGF2/α1-subCD3 binding interface by in silico docking studies and atomistic molecular dynamics simulations

To analyze the protein-protein interface between FGF2 and the cytoplasmic domain of Na,K-ATPase and to evaluate a potential role of K54 and K60 as residues in FGF2 that are in direct contact with α1-subunit, we conducted both *in silico* molecular docking studies and atomistic molecular dynamics (MD) simulations. In a first stage, using protein-protein docking protocols, we scanned possible interaction interfaces between α1-subCD3 and the region in FGF2 exposing K54 and K60. Intensive molecular docking simulations were conducted by rotating FGF2 around the α1-subCD3 domain and calculating the corresponding interface scores, i.e. the differences in the energy state between the FGF2/α1-subCD3 complex versus unbound FGF2 and unbound α1-subCD3. The docking results were filtered and clustered based upon the root-mean-square deviation (RMSD) for FGF2 and the most representative structures of the largest populated clusters were refined employing MD simulations as explained in Material and Methods. The ultimate goal of this approach was to find the most stable FGF2/α1-subCD3 interface and to characterize the network of interactions it involves. In Fig. 8D, the position of FGF2 relative to the full-length Na,K-ATPase is illustrated for the WT1 cluster. The human version of the α1-subCD3 domain was aligned to the same domain of the full–length crystal structure of the α1-subunit of the Na,K-ATPase from *Sus scrofa* (Residues T380 – V597; PDB ID: 3KDP). Membrane lipids [phosphatidylcholine and PI(4,5)P_2_] were added to visualize the position of FGF2 bound to α1 relative to the inner leaflet of the plasma membrane. To test the stability of the interface between FGF2 and α1-subCD3, the central structure [which represents the structure with the smallest average value for the root-mean-square deviation (RMSD) compared to all other structures from the same cluster] of the cluster was simulated in three replicates for 200 ns. The probabilities of physical contacts as average contact maps of the FGF2/α1-subCD3 interface reveals a broad network of interactions stabilizing the interface of the WT1 system (Fig. 8A). As shown in the interaction energies map (Fig. 8C), the FGF2/α1-subCD3 interface is strongly stabilized by three ion pairs, E525– K60, E498–R30 and K524–D27. Furthermore, K54 was found to be important as it stabilized the interaction between E498 and R30. The contribution of each residue in stabilizing the FGF2/α1-subCD3 interface (Fig. 8B), calculated by summing up the probabilities of contact for each residue individually, indicates that K54 and K60 have a high probability to be involved in the interaction.

**Fig. 8:**
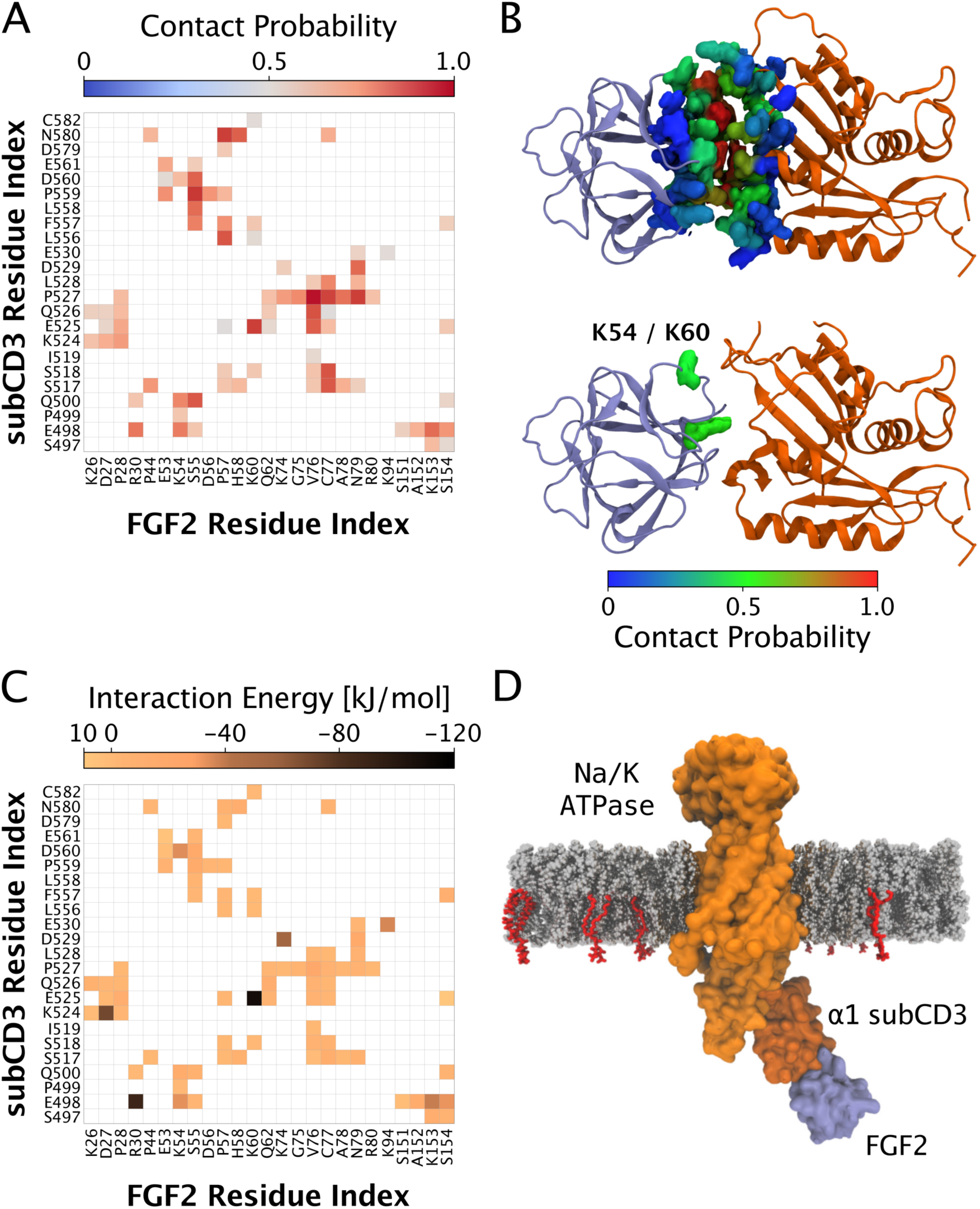
Characterization of the molecular interface between FGF2 and α1-subCD3 based on in silico docking studies and atomistic molecular dynamics simulations. In panel A, a pairwise contact map with all residues characterized by a probability of contact of more than 50% is shown for the WT1 simulated system. In panel B, an average structure of the FGF2/α1-subCD3 interface is shown illustrating the contribution to the interaction for each residue individually. It is defined as the sum of the probabilities of contacts for each residue and it is represented as a colored surface using the RGB color scale. In panel C, the average interaction energy (electrostatic and van der Waals contributions) of each FGF2/α1-subCD3 pair with a probability of contact >50% is shown. In panel D, the most representative structure of the WT1 cluster is illustrated. The human α1-subCD3 domain was aligned to residues T380–V597 of the crystal structure of the α1-subunit of the Na,K-ATPase from *Sus scrofa* (PDB ID: 3KDP). The Na,K-ATPase is represented as an orange surface with the α1-subCD3 domain highlighted using a darker shade. FGF2 is shown as a violet surface. The PI(4,5)P_2_ and phosphatidylcholine membrane lipids are represented using van der Waals spheres with red and grey colors, respectively.

In conclusion, the *in silico* results shown in Fig. 8 suggest the WT1 structure (representing the first cluster) to be the most probable protein-protein interaction surface between FGF2 and α1-subCD3. It depends on K54 and K60 and is characterized by a large number of stable contacts with three ion pairs that are stable throughout the MD simulation time. Furthermore, MD simulations comparing FGF2-wt with FGF2-K54/60E provided direct evidence for the relevance of the WT1 system as the relevant interface between FGF2 and α1-subCD3. This is evident from the observation that, in all MD simulations of the mutated system (for details see Materials and Methods) and as opposed to FGF2-wt, FGF2-K54/60E did either dissociate from or was not able to form a stable interaction with α1-subCD3 (Video S1). Thus, the molecular docking studies and MD simulations presented in Fig. 8 as well as in Video S1 are consistent with the experimental data shown in Figs. 1 to 7.

### The α1 subunit of the Na,K-ATPase contributes to FGF2-GFP recruitment at the inner leaflet of the plasma membrane

Using a single particle TIRF microscopy approach established previously ^12^, we quantified FGF2-GFP recruitment at the inner plasma membrane leaflet of living cells. For the wild-type and the various mutant forms of FGF2 indicated, both widefield and TIRF images were taken (Fig. 9). While the widefield images allowed for the analysis of total expression levels of each of the FGF2-GFP fusion proteins indicated, the TIRF images were processed for the quantification of individual FGF2-GFP particles per surface area in the vicinity of the plasma membrane. In panel A, FGF2-GFP fusion proteins are shown in which K54 and K60 were substituted for glutamates and compared with FGF2 wild-type. The results shown in Fig. 9A along with those from real-time videos monitoring FGF2-GFP recruitment (Video S2) suggested that FGF2-K54/60E is impaired regarding physical contacts with the inner leaflet of the plasma membrane. A larger data set for each of the FGF2 mutants shown in Fig. 9A and Video S2 was subjected to quantification and a statistical analysis as shown in Fig. 10. The number of FGF2-GFP particles in the vicinity of the plasma membrane were normalized to surface area with each data point representing the analysis of one cell. These experiments demonstrated that substitution of K54 and K60 by glutamates reduced the number of FGF2-GFP particles at the inner plasma membrane leaflet by about 50% (Fig. 10A and 10C). In additional experiments, the substitution of K54 and K60 was combined with mutations in the PI(4,5)P_2_ binding site of FGF2 (Fig. 9B, 10B, 10C and Video S3). These experiments revealed that mutations restricted to the PI(4,5)P_2_ binding site in FGF2 alone did not affect FGF2-GFP recruitment at the inner plasma membrane leaflet (Fig. 10B and 10C). In addition, the combination of substitutions of K54/K60 and mutations in the PI(4,5)P_2_ binding site of FGF2 caused a similar phenotype compared to what was observed for K54/60E alone (Fig. 10B and 10C). The differences in the efficiency of FGF2 recruitment at the inner leaflet between FGF2 mutants defective in either α1 binding (FGF2-K54/60E) or PI(4,5)P_2_binding (FGF2-K127Q/R128Q/K133Q) are particularly evident following GFP background correction (Fig. 10C). While substitution of K54 and K60 by glutamates reduced FGF2-GFP recruitment by almost 50%, the FGF2 mutant incapable of binding to PI(4,5)P_2_ was fully functional with regard to recruitment at the inner plasma membrane leaflet. In addition, when both sets of amino acid substitutions were combined, the observed phenotype was not stronger compared to the K54/60E substitution alone (Fig. 10C). These findings suggest that, during recruitment at the inner plasma membrane leaflet, the physical contact of FGF2 with α1 subunit of the Na,K-ATPase precedes the interaction of FGF2 with PI(4,5)P_2_. Furthermore, in a cellular context, the data shown in Figs. 9 and 10 imply that the interaction of FGF2 with α1 facilitates the subsequent binding of FGF2 to PI(4,5)P_2_, i.e. α1 acts upstream of PI(4,5)P_2_.

**Fig. 9:**
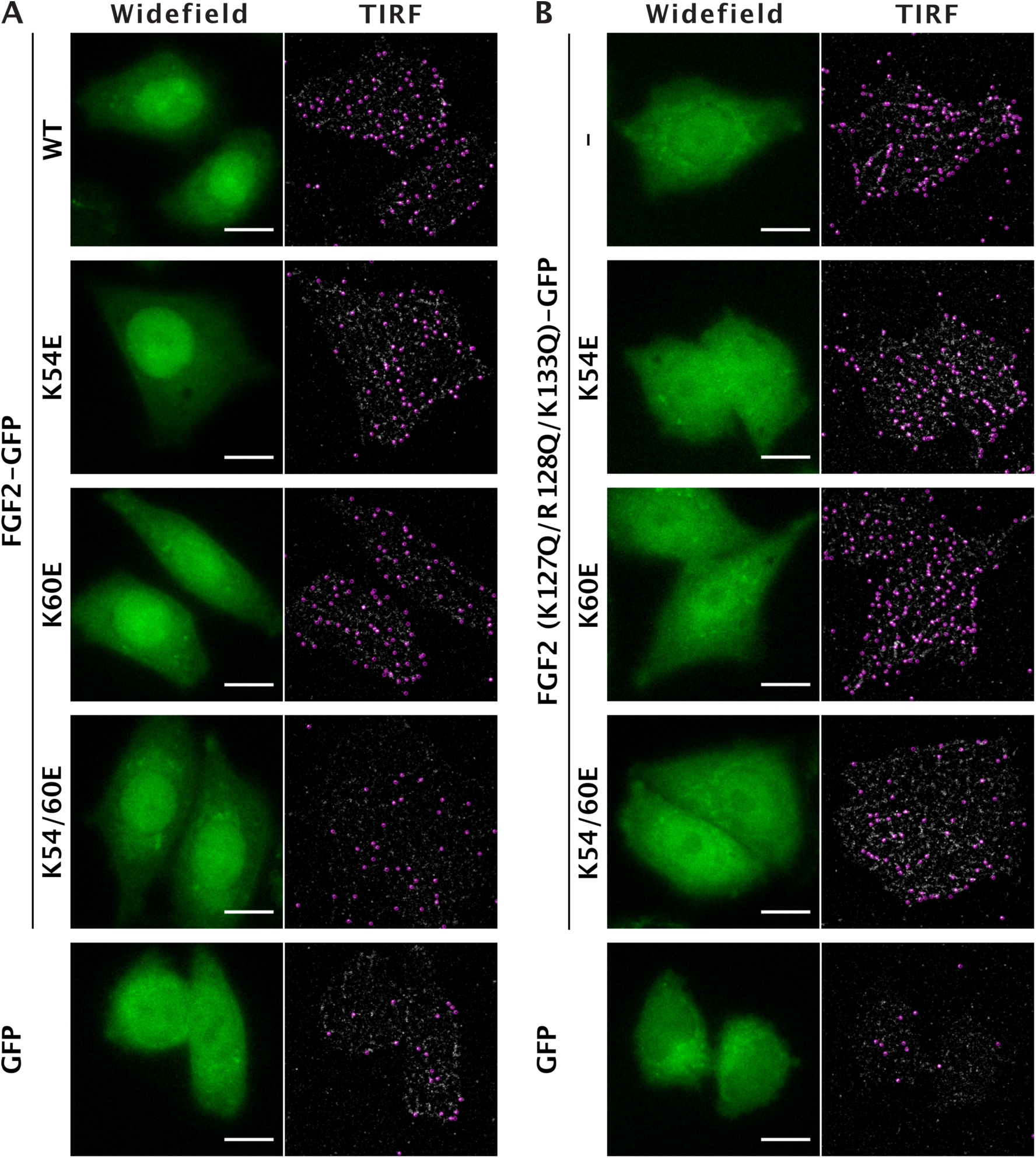
Quantitative analysis of FGF2-GFP membrane recruitment in living cells at the single particle level. A) Stable CHO-K1 cell lines expressing either FGF2-wt-GFP, FGF2-K54E-GFP, FGF2-K60E-GFP, FGF2-K54/60E-GFP or GFP in a doxycycline-dependent manner were imaged employing high resolution TIRF microscopy as reported previously^12^. Single FGF2-GFP or GFP particles were identified at the inner plasma membrane leaflet (labeled by pink circles). For each cell line, a widefield image and the first frame of the corresponding TIRF video are shown. Scale bar = 10 μm. B) Stable CHO-K1 cell lines expressing either FGF2-wt-GFP, FGF2-K127Q/R128Q/K133Q-GFP, FGF2-K54E/K127Q/R128Q/K133Q-GFP, FGF2-K60E/K127Q/R128Q/K133Q-GFP, FGF2-K54/60E/K127Q/R128Q/K133Q-GFP or GFP in a doxycycline-dependent manner were cultured and imaged as described in panel A.

**Fig. 10:**
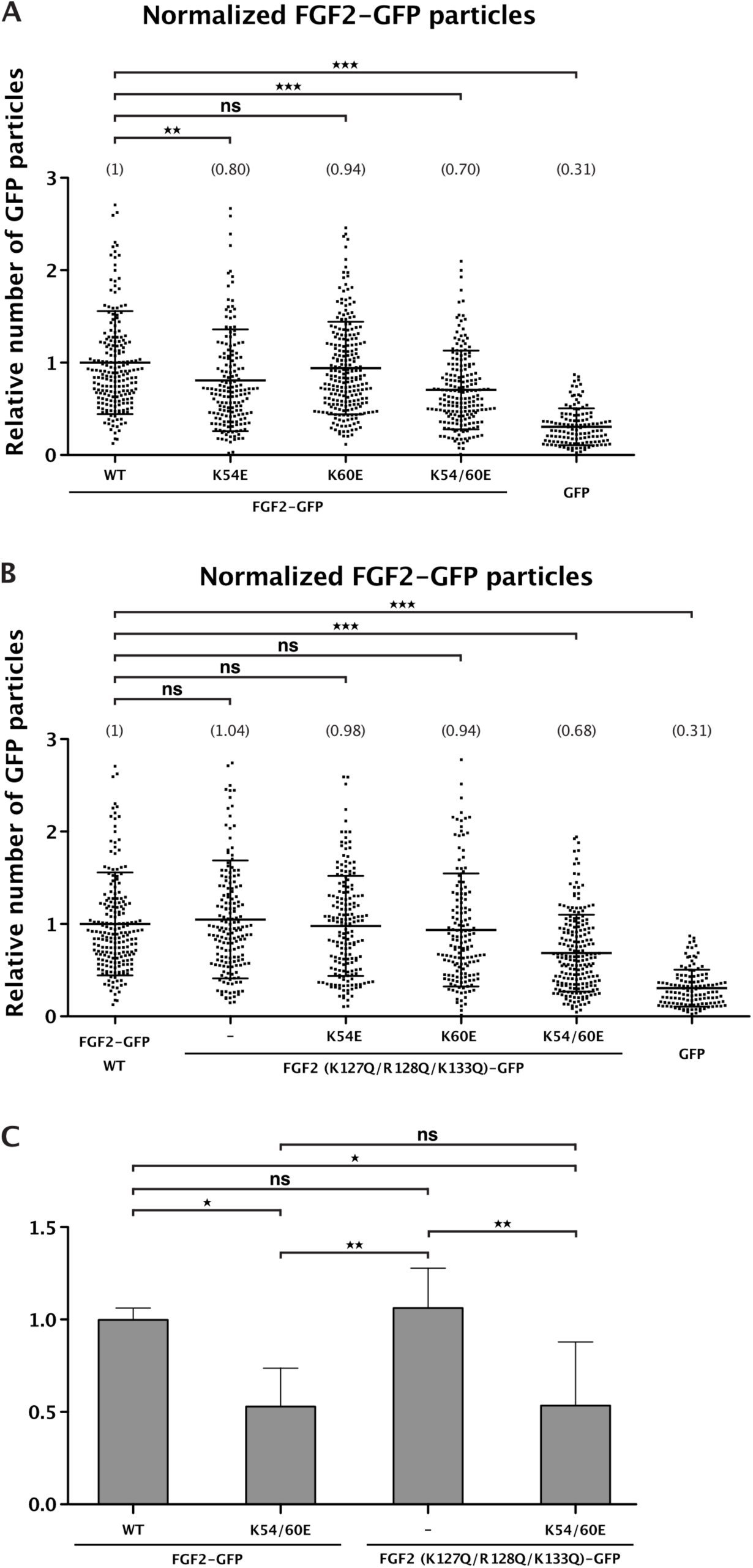
FGF2-GFP recruitment at the inner leaflet depends on direct interactions with the cytoplasmic domain of α1. A) FGF2-wt versus FGF2-K54/60E (FGF2 mutant deficient in binding to α1) B) FGF2-wt versus FGF2-K54/60E in a K127Q/R128Q/K133Q background (FGF2 mutant deficient in binding to PI(4,5)P_2_) C) Direct comparison between FGF2-K54/60E and FGF2-K127Q/R128Q/K133Q following GFP background subtraction Quantification of FGF2-GFP membrane recruitment at the inner leaflet of intact cells for all wild-type and mutant forms of FGF2 shown in panels A and B of Fig. 10. Time-lapse TIRF videos with a total of 100 frames (100 ms/frame) were analyzed using the Fiji plugin TrackMate ^12^. A total of four biological replicates were included in the analyses shown in panels A and B. The number of GFP particles were normalized for both surface area and the relative expression levels of each FGF2 fusion protein in the corresponding cell line. In panel A, FGF2 mutants defective in binding to α1 are shown (K54E, K60E and K54/60E). In panel B, the same mutants were combined with mutations in the PI(4,5)P_2_ binding pocket of FGF2 (K127Q/R128Q/K133Q). The mean values of each condition are shown in brackets with the wild-type form of FGF2-GFP set to 1. In panel C, the most important conditions were directly compared following GFP background subtraction and are shown as bar graphs. The mean values for FGF2-wt-GFP, FGF2-K54/60E-GFP, FGF2-K127Q/R128Q/K133Q-GFP and FGF2-K54/60E/K127Q/R128Q/K133Q-GFP are given along with a statistical analysis using six biological replicates. All statistical analyses were based on a one-way ANOVA test combined with Tukey’s post hoc test (ns = p ≥ 0.05; *** = p ≤ 0.001).

### Unconventional secretion of FGF2-GFP is impaired upon substitution of K54 and K60 by glutamates

Beyond quantifying the recruitment of FGF2-GFP at the inner leaflet in living cells (Figs. 10 and 11), we tested whether amino acid substitutions that impair the interaction of FGF2 with α1 subunit of the Na,K-ATPase do affect secretion of FGF2 to the cell surface (Fig. 11). These experiments included various FGF2 mutants in which K54 and K60 were replaced by glutamates (Fig. 11A and 11B) and were also combined with amino acid substitutions (K127Q/R128Q/K133Q) that compromise the ability of FGF2 to bind to PI(4,5)P_2_ (Fig. 11C and 12D). Following induction of protein expression with doxycycline, a well-established cell surface biotinylation assay was used to quantify the extracellular amounts of the FGF2-GFP fusion proteins indicated ^20, 23, 41, 43^. These experiments revealed that a K54/60E substitution alone caused a significant impairment of FGF2-GFP transport to the cell surface with only 60% efficiency compared to the wild-type form of FGF2-GFP (Fig. 11A and 11B). When K54/60E and K127Q/R128Q/K133Q substitutions were combined, the efficiency of FGF2-GFP secretion dropped to less than 20% compared to the wild-type form of FGF2-GFP (Fig. 11C and 11D). This phenotype was similar to a mutant form of FGF2-GFP that lacks two cysteine residues (C77/95A) required for FGF2 oligomerization and the formation of membrane translocation intermediates, the so far most severely affected FGF2 secretion mutant known in the literature ^22, 23^. Our combined findings suggest that efficient secretion of FGF2-GFP into the extracellular space is facilitated by a direct interaction of FGF2 with the cytoplasmic domain of α1 subunit of the Na,K-ATPase, a physical contact that precedes the interaction of FGF2 with PI(4,5)P_2_.

**Fig. 11:**
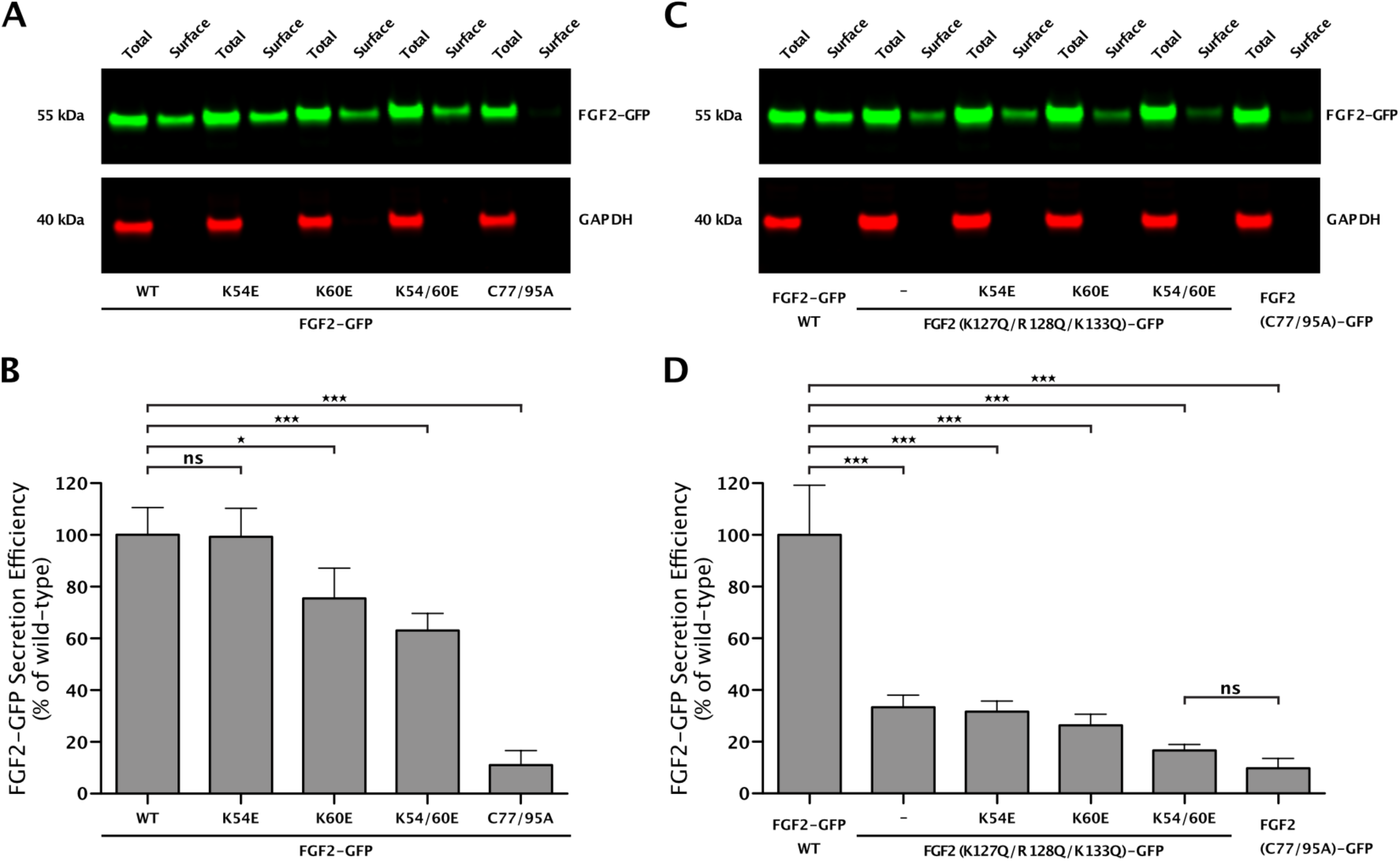
Efficient secretion of FGF2 from cells is facilitated by its interaction with α1-subunit of the Na,K-ATPase. A) Cell surface biotinylation experiments were conducted as described in Materials and Methods using stable CHO-K1 cell lines expressing either FGF2-wt-GFP, FGF2-K54E-GFP, FGF2-K60E-GFP, FGF2-K54/60E-GFP or FGF2-C77/95A-GFP in a doxycycline-dependent manner. Aliquots from the total cell lysate (1.6%) and from the biotinylated fraction (33.3%; corresponding to the cell surface population of proteins) were subjected to SDS-PAGE and Western blotting. Anti-GFP antibodies were used to detect the various FGF2-GFP fusion proteins indicated. Anti-GAPDH antibodies were used to detect intracellular GAP-DH as a control for cell integrity during cell surface biotinylation. Primary antibodies were detected by fluorophore-labeled secondary antibodies and quantified using the Odyssey^®^ CLx Imaging System (LI-COR Biosciences). B) The efficiency of FGF2-GFP secretion of each variant form shown in panel A was quantified and normalized to the wild type form that was set at 100%. The statistical analysis was based on a one-way ANOVA test combined with Tukey’s post hoc test (ns = p ≥ 0.05; *** = p ≤ 0.001). C) Stable CHO-K1 cell lines expressing either FGF2-wt-GFP, FGF2-K127Q/R128Q/K133Q-GFP, FGF2-K54E/K127Q/R128Q/K133Q-GFP, FGF2-K60E/K127Q/R128Q/K133Q-GFP, FGF2-K54/60E/K127Q/R128Q/K133Q-GFP or FGF2-C77/95A-GFP in a doxycycline-dependent manner were analyze by cell surface biotinylation as described in the legend to panel A. D) The efficiency of secretion of each variant form of FGF2-GFP shown in panel C was quantified and normalized as described in the legend to panel B. The statistical analysis was based on a one-way ANOVA test combined with Tukey’s post hoc test (ns = p ≥ 0.05; *** = p ≤ 0.001).

## Discussion

FGF2 is a cell survival factor with a critical role in tumor-induced angiogenesis ^8, 9^. However, unlike the majority of extracellular factors including functionally related proteins such as VEGF, FGF2 is secreted by an unconventional secretory pathway that is mechanistically distinct from the classical ER/Golgi-dependent pathway of protein secretion in eukaryotic cells ^5–7^. Unconventional secretion of FGF2 is mediated by direct translocation across the plasma membrane and results in transport of FGF2 to the cell surface where it remains bound to heparan sulfate chains of proteoglycans ^13, 14, 20, 21, 38, 39^. Thence, FGF2 can be exchanged between heparan sulfate chains on cell surfaces to form ternary signaling complexes with FGF receptors for both autocrine and paracrine signaling ^8, 9^.

The core mechanism of FGF2 membrane translocation has been investigated in great detail. This includes the central role of the phosphoinositide PI(4,5)P_2_ in (i) FGF2 membrane recruitment at the inner leaflet, (ii) the insertion of membrane-spanning FGF2 oligomers that serve as translocation intermediates and (iii) the formation of a lipidic membrane pore with a toroidal architecture ^5, 7, 10, 12, 17–19, 22, 23, 44–46^. In addition, the role of cell surface heparan sulfates in mediating the final step of FGF2 membrane translocation is well understood ^5, 12, 20–22, 38, 39, 44^. By contrast, the functional role of α1 subunit of the Na,K-ATPase in unconventional secretion of FGF2 is unknown ^5, 7^.

Initial evidence for a role of α1 in FGF2 secretion came from pharmacological studies. It was found that ouabain, a well-established inhibitor of the Na,K-ATPase, blocks FGF2 secretion from fibroblasts ^33^. These studies were confirmed in other cell types ^35^ and were further corroborated by experiments demonstrating a rescue of FGF2 secretion in the presence of ouabain, provided an α1 variant form was expressed to which ouabain cannot bind ^36^. The pharmacological studies with ouabain were then confirmed by technically independent approaches demonstrating a reduced efficiency of FGF2 secretion upon RNAi mediated down-regulation of α1 ^15^. Furthermore, a direct interaction of FGF2 with the cytoplasmic domain of α1 (K_D_: ∼0.8 µM) was reported ^15^. However, while a role for α1 in unconventional secretion of FGF2 is an accepted fact in the field, its functional role in this process is unknown.

In the current study, we identified a sub-domain in the cytoplasmic part of α1 (α1-subCD3) that mediates a direct physical interaction with FGF2. α1-subCD3 belongs to the nucleotide-binding domain of α1. The basic unit of this interaction is a heterodimer with a K_D_ in the sub-micromolar range. The identification of α1-subCD3 as a minimal binding partner of FGF2 with about 20 kDa in size allowed for solution NMR experiments. This led to the identification of two lysine residues in position 54 and 60 on the molecular surface of FGF2 that were found critical for the interaction with α1-subCD3. Intriguingly, even though FGF family members in general are highly conserved reflecting the structural needs for building up the typical FGF fold, K54 and K60 represent FGF2-specific residues that are absent from most FGF family members carrying signal peptides for ER/Golgi dependent protein secretion. Therefore, similar to what has previously been reported for two cysteine residues on the molecular surface of FGF2 ^23^, the exclusive presence of both K54 and K60 in FGF2 points at a specific function of these residues in unconventional secretion of FGF2. Using various kinds of protein-protein interaction assays, we confirmed the NMR results in that mutants of FGF2 lacking K54/K60 residues were impaired in binding to α1-subCD3. These experiments were further validated by *in silico* docking studies and atomistic molecular dynamics simulations demonstrating a role for K54/K60 in FGF2 binding to α1-subCD3 in a thermodynamically relevant model system. These findings were further found to be functionally relevant in a cell-based model system with FGF2 secretion being impaired in the absence of K54/K60. Finally, we made use of a recently established single particle imaging system studying FGF2 membrane recruitment at the inner plasma membrane leaflet employing real-time TIRF microscopy in living cells ^12^. Intriguingly, we found that a FGF2 mutant lacking both K54 and K60 as well as the three critical residues (K127/R128/K133) of the PI(4,5)P_2_ binding pocket was indeed impaired in binding to the inner leaflet. However, when the two sets of mutations were looked at separately, only the removal of K54/K60 caused a decrease in FGF2 recruitment at the inner leaflet. These findings have two important implications with (i) FGF2 binding to the α1-subunit of the Na,K-ATPase precedes FGF2 binding to PI(4,5)P_2_ and (ii) the physical contact of FGF2 at the inner leaflet with α1 facilitates subsequent interactions of FGF2 with PI(4,5)P_2_. Thus, beyond the identification of both a sub-domain in the cytoplasmic part of α1 and residues on the surface of FGF2 required for a direct interaction between these proteins, our findings suggest a function of α1 as an auxiliary factor in unconventional secretion of FGF2 that increases the efficiency of PI(4,5)P_2_-dependent FGF2 oligomerization and membrane translocation. In this context, it is an interesting observation that, as opposed to *in vitro* experiments in which FGF2 can bind to PI(4,5)P_2_ in the absence of other factors ^18, 19^, α1 appears to be important in intact cells to enable efficient binding of FGF2 to PI(4,5)P_2_. Thus, the α1-subunit of the Na,K-ATPase may serve as a factor that accumulates FGF2 at the inner plasma membrane leaflet, facilitating PI(4,5)P_2_ dependent FGF2 membrane translocation to the cell surface. Furthermore, it is an interesting hypothesis for future studies as to whether this function is linked to a modulation of the activity of the Na,K-ATPase to maintain a functional Na/K gradient across the plasma membrane during active events of FGF2 membrane translocation, a process that may transiently disturb the integrity of the plasma membrane.

Recently, beyond the established function of α1 in unconventional secretion of FGF2, evidence has been reported for a similar role of α1 in the secretion of HIV-Tat from HIV-infected T cells ^37^. Along with the well-established role for PI(4,5)P_2_ in unconventional secretion of HIV-Tat ^30, 31^, these findings point at a shared secretory mechanism between FGF2 and HIV-Tat. Furthermore, with Tau and Interleukin 1β, two additional extracellular factors secreted by unconventional means, even more examples have been reported for unconventional secretory processes that depend on PI(4,5)P_2_ as a prerequisite for their transport into the extracellular space ^27–29, 32^. Thus, after decades of research on unconventional secretory mechanisms in mammalian cells, common features between pathways that are taken by proteins functionally as different as FGF2, Interleukin 1β, Tau and HIV-Tat are beginning to emerge.

## Acknowledgements

This work was funded by the Deutsche Forschungsgemeinschaft (DFG, German Research Foundation) - Project Number 278001972 - TRR 186 and Ni 423/7-1. This work was further supported by the DFG Cluster of Excellence CellNetworks at Heidelberg University, the Academy of Finland (Center of Excellence program), and the Sigrid Juselius Foundation. We thank Monika Langlotz from the ZMBH Flow Cytometry Core Facility (Zentrum für Molekulare Biologie Heidelberg) and Holger Lorenz from the ZMBH Light Microscopy Core Facility (Zentrum für Molekulare Biologie Heidelberg) for their support in cell sorting and image acquisition/interpretation, respectively. C.F. and J.S. would like to acknowledge the assistance of the Core Facility BioSupraMol at Freie Universität Berlin, which is supported by the DFG. For computational resources, we wish to thank the CSC – IT Center for Science (Espoo, Finland). F.L. was supported by a HPC-Europa3 Transnational Access program (HPC175W35X).

## Materials and Methods

### Recombinant proteins and biochemical protein-protein interaction assays

Recombinant proteins were expressed in *E. coli* and purified according to standard procedures. The 18 kDa isoform of FGF2 was expressed and purified as a N-terminally His_6_-tagged protein. For competition assays, a non-tagged form of FGF2 lacking 25 residues at the N-terminus (ΔN25-FGF2) was expressed and purified. For NMR spectroscopy, a monomeric FGF2 variant form (C77/95S) was expressed in M9 minimal medium with ^15^N-NH_4_Cl as the sole nitrogen source in order to produce FGF2 as a ^15^N labeled protein. In addition, recombinant forms of FGF2 were expressed and purified in which K54 and K60 were substituted by glutamates as indicated.

Four variant forms of the cytoplasmic domain of α1 were expressed as N-terminal GST fusion proteins. This included a fusion of the three main loops of the cytoplasmic domain [GST-α1-CD1-3; ^15^], the third loop of the cytoplasmic domain alone [GST-α1-CD3; C343-L772 from human α1); ^15^], a small sub-domain of loop 3 (GST-α1-subCD3; T382–A598 from human α1) and the third cytoplasmic loop of α1 lacking the above mentioned sub-domain of loop 3 (GST-α1-CD3Δsub; C343-L381 linked to A598-L772 from human α1).

### Biochemical pull-down experiments to analyze interactions between FGF2 and α1

For this set of experiments, Glutathione Sepharose beads were equilibrated in PBS buffer and incubated with the GST-tagged variants of α1 as indicated. GST alone was used as a negative control. GST-coupled Sepharose beads were blocked with 3% (w/v) BSA in PBS supplemented with 1 mM Benzamidine and 0.05% (w/v) Tween 20 (buffer A), washed extensively in buffer A. Finally, beads were resuspended with five bed volumes of buffer A. Per experimental condition, 75 µl of beads solution were incubated with 15 µg of His-tagged FGF2 in a total volume of 200 µl of buffer A for one hour at room temperature. Following collection of beads by low-speed centrifugation and extensive washing with buffer A, bound protein was eluted with SDS sample buffer. Both bound (33%) and unbound (2.5%) material was analyzed by SDS-PAGE followed by protein staining using Coomassie InstantBlue. Gels were scanned using the Odyssey^®^ CLx Imaging System (LI-COR Biosciences) and band intensities were quantified using the LI-COR ImageStudio software. For quantification, the ratios between bound and unbound material were calculated for each experimental condition as indicated.

### Quantitative analysis of the interaction between FGF2 and α1 employing AlphaScreen technology

For cross-titration of a wide range of concentrations of FGF2 and α1 variants and to determine affinity between FGF2 and α1 the AlphaScreen® protein-protein interaction assay was used [Figs. 2 and 7; ^15^]. N-terminally His-tagged FGF2 (wild-type or mutants as indicated) and N-terminally GST-tagged α1 variants were cross-titrated from 500nM to 8nM in PBS supplemented with 0.1% (w/v) BSA and 0.05% (w/v) Tween 20. These experiments were conducted in 384-well plates. After one hour of incubation, the proteins were incubated with AlphaScreen Ni-NTA acceptor and AlphaScreen Glutathione donor beads, each at a final concentration of 11.25 µg/ml in a total volume of 15 µl. Following two additional hours of incubation, samples were measured using an EnVision plate reader (PerkinElmer). For each biological replicate, data were normalized to the signal measured for His-FGF2/GST-α1 CD1-3 (Fig. 2) or for His-FGF2 WT/GST-α1-subCD3 (Fig. 7) at 250nM FGF2 and 63nM GST-tagged protein. Data were represented as a heat map with the highest signal set to 100% (displayed in red) and the lowest signal set to 0% (displayed in blue) as indicated by the color legend.

To determine binding affinity between His-FGF2-wt and GST-α1 variants as indicated, competition experiments were conducted. An untagged and N-terminally truncated form of FGF2 (ΔN25-FGF2) was used as a competitor which allowed for the determination of IC50 values. For these analyses, optimal concentrations of FGF2 (62 nM) and α1 (15 nM) were used based upon the cross-titration experiments described above. The competitor ΔN25-FGF2 was used in a concentration range between 22.5 µM and 1.4 nM. The respective pairs of His-and GST-tagged proteins were mixed with the ΔN25-FGF2 competitor in a final volume of 10 µl at the concentrations indicated. Addition of Alpha beads and measurements were done as described above. For each protein pair, the median signal of three technical replicates was calculated, normalized to the signal of the buffer lane and plotted against the concentration of the ΔN25-FGF2 competitor. The error estimates are based on three independent biological replicates each of which consisted of three technical replicates. The competitor concentration promoting half-maximal inhibition (IC50) of the signal was determined by fitting the experimental data with a non-linear regression model [log(inhibitor) vs. response - variable slope (four parameters)] using GraphPad Prism 5.0c software. Under suitable experimental conditions, the apparent IC50 value in this type of competition experiments corresponds to the dissociation constant of the observed protein-protein interaction.

### Analysis of FGF2 variant forms regarding PI(4,5)P2-dependent oligomerization, binding to heparin and thermal stability

To test the ability of FGF2 variant forms to bind to PI(4,5)P_2_ concomitant with oligomerization, FGF2 (10µM) was incubated with liposomes (2 mM total lipid) with a plasma-membrane-like lipid composition containing 3 mol% of PI(4,5)P_2_. As described previously ^17, 18^, experiments were conducted in a final volume of 50 µl HK-buffer (150 mM KCl, 25 mM HEPES, pH 7.4) in an Eppendorf Thermomixer at 25°C at 500rpm. After 4 hours of incubation liposomes were sedimented for 10 min at 16.000g, and washed with 100µl HK-buffer. Pellets were dissolved in 20 µl non-reducing SDS sample buffer and heated for 10 min at 65°C. Samples were analyzed on 1.0 mm NuPAGE 4-12% non-reducing Bis-Tris gels/MES-running buffer (Invitrogen) and proteins were stained with Coomassie InstantBlue (Expedeon). Input lanes contained 50% of FGF2 used in the oligomerization assays.

FGF2 variant forms were further compared to FGF2-wt with regard to their ability to bind to heparin. Heparin Sepharose 6 Fast Flow (GE Healthcare) beads (10 µl slurry) were washed four times in PBS and incubated for 1 hour at room temperature with the FGF2 variant forms indicated (15µg protein each in a total volume of 200 µl PBS). Following sedimentation of the beads and extensive washing, binding efficiency was analyzed by SDS-PAGE and protein staining using Coomassie InstantBlue (Expedeon).

To determine native protein folding by means of thermal stability, the various FGF2 forms used in this study were analyzed in nanoDSF experiments using a Prometheus NT 48 instrument (Nanotemper) ^47^. This procedure monitors the light absorbance of proteins at 330nm and 350nm along a thermal gradient. The ratio of A_330nm_/A_350nm_ was plotted as a function of the temperature. This allows for determining the unfolding transition midpoint, i.e. the melting temperature of a protein. The FGF2 variant forms indicated were used in a volume of 10 µl at a final concentration of 1.5 mg/ml.

### Crosslinking experiments

The crosslinking experiments shown in Fig. 3 were performed at 25°C in 25 µl HK buffer (25 mM Hepes, pH 7.4; 150 mM KCl) at a final protein concentration of 10 µM. When FGF2 and α1-subCD3 were mixed, the final protein concentration was 20 µM, irrespective of the ratio between FGF2 and α1-subCD3 (1:1, 1:2 and 2:1). As chemical crosslinkers, BMOE (bismaleidomethane) and DSG (disuccinimidyl glutarate) were dissolved in 20 mM DMSO and further diluted in HK buffer to 0.5 mM (BMOE) and 2 mM (DSG), respectively. Following preincubation of proteins for 30 min, reactions were started with the addition of BMOE or DSG yielding crosslinker/protein ratios of 1:1 (BMOE) and 4:1 (DSG). After 30 min, samples were quenched with 10 mM DTT (BMOE) or 20mM Tris/HCl pH 7.5 (DSG), mixed with an equal volume of SDS sample buffer containing β-mercaptoethanol and incubated for 10 min at 70°C. Of each sample, 80% were analyzed on 1.5 mm NuPAGE 4-12% Bis-Tris gels (Invitrogen) and stained with Coomassie (InstantBlue, Expedeon). For a Western analysis, 2% of each sample were separated on 1 mm NuPAGE 4-12% Bis-Tris gels. Blots were analyzed using affinity-purified anti-α1-CD1-3 rabbit antibodies ^15^ and monoclonal anti-FGF2 antibodies (clone bFM-1, Millipore).

### Structural analysis of the FGF2/α1-subCD3 interface employing NMR spectroscopy

NMR spectra were recorded at 300K on a Bruker Avance III 700MHz spectrometer equipped with a 5mm triple resonance cryo-probe. ^1^H-^15^N-HSQC spectra were acquired of 77 µM ^15^N-labeled FGF2-C77/95S in the absence or presence of 70 µM or 138 µM α1-subCD3 in 25 mM HEPES (pH 7.4), 150mM KCl, 10% D_2_O with 108 scans and 1024 data points in the ^1^H and 96 data points in the ^15^N dimension. Spectra were processed with Topspin3.2 (Bruker, Billerica, USA) and analyzed using CcpNMR Analysis 2.4.2 ^48^. Peak assignments were transferred from published data [BMRB entry 4091; ^42^] to nearest neighbors in recorded spectra, if possible, leading to an assignment of 63% of the non-Proline residues. Chemical shift differences were calculated using the formula Δδ(^1^H,^15^N)=(δ(^1^H)^2^+(0.15*δ(^15^N))^2^)^1/2^. Signal-to-noise ratios were calculated from peak intensities and average noise levels. Overlapping peaks were omitted from the analysis.

### Multiple sequence alignment performed with FGF family members

In order to identify residues that uniquely present in FGF2 compared to signal-peptide-containing FGF family members, a multiple sequence alignment was performed using the EMBL-EBI MUSCLE 3.8 tool ^49^. The protein sequence of isoform 3 of FGF2 (18 kDa form; UniprotKB ID P09038-2) was aligned with canonical sequences of FGF1 (P05230), FGF3 (P11487), FGF4 (P08620), FGF5 (P12034), FGF6 (P10767), FGF7 (P21781), FGF8 (P55075), FGF9 (P31371), FGF10 (O15520), FGF11 (Q92914), FGF12 (P61328), FGF13 (Q92913), FGF14 (Q92915), FGF16 (O43320), FGF17 (O60258), FGF18 (O76093), FGF19 (O95750), FGF20 (Q9NP95), FGF21 (Q9NSA1), FGF22 (Q9HCT0) and FGF23 (Q9GZV9). The alignment covered residues 32 to 105 (FGF2).

### In silico docking studies

The human sequence of the α1-subCD3 domain (T382–V599) was modeled based upon the existing crystal structure of the Na,K-ATPase of *Sus scrofa* [Residues T380-V597, PDB ID: 3KDP; ^50^]. The two structures differ in only 6 amino acids, which is less than 3% of the total amino acid content. Point mutations using the CHARMM-GUI web server ^51^ were employed to model the human sequence of the α1-subCD3 domain as described in Table 1.

**Table 1.** Point mutations carried out to model the human sequence of the α1 subCD3 domain, starting from the crystal structure of the Na/K ATPase of *Sus scrofa*.

| Residue ID | Amino Acid | Mutation |
| --- | --- | --- |
| 391 | S | F |
| 473 | T | A |
| 497 | A | S |
| 500 | R | Q |
| 521 | I | L |
| 578 | L | I |

Protein-protein docking studies were performed using the Rosetta 2018 package ^52–55^. The α1-subCD3 domain was first considered as a spherical unit, and FGF2 was rotated around this sphere, positioning the surface area of FGF2 that contains K54 and K60 towards the α1-subCD3 surface. About 120,000 structures were generated with the Rosetta global docking protocol. About 94% of the structures were discarded based on low interface scores. All structures without contacts between K54 and K60 of FGF2 and the α1-subCD3 domain were discarded as well. Here, a contact was defined if the distance between any atoms of K54 or K60 in FGF2 and α1-subCD3 was less than 6 Å. The remaining structures were aligned to the α1-subCD3 domain of the full-length *Sus scrofa* crystal structure of Na,K-ATPase. All docked structures with overlaps of FGF2 and α1-subCD3 domains were also removed. This filtering procedure resulted in 62 candidates of docked structures for further analysis. For each of them, the Rosetta local docking protocol was employed to refine the global docking results. Therefore, for each of the 62 candidates, 500 structures were generated by randomly perturbing FGF2 by 3 Å translation and 8° rotation before the start of every individual simulation. These 31,000 structures were subjected to the same filtering procedure, which reduced the set to 33 structures. These were clustered based on the RMSD value for FGF2 using the Gromos algorithm ^56^ and a RMSD cut-off for two structures to be neighbors within 0.6 nm, which in turn resulted in four most populated clusters C1-C4 (Table 2). In each cluster, the most representative structure (as the centroid of the structures in a given cluster) was chosen as the basis to represent the wild-type (WT1-WT4), and the FGF2-α1-subCD3 interfaces of these structures were then tested and refined employing atomistic molecular dynamics simulations.

**Table 2.** Number of structures and the averaged root-mean-square deviation (RMSD) values for the four most popular clusters identified in the docking analyses.

| Cluster | Number of Structures | <RMSD> (nm) |
| --- | --- | --- |
| C1 | 11 | 0.232 |
| C2 | 4 | 0.146 |
| C3 | 4 | 0.252 |
| C4 | 4 | 0.330 |

### Atomistic molecular dynamics simulations

The screening of the most probable FGF2/α1-subCD3 interaction interface was conducted through an in-depth study of the type of molecular interactions taking place during the simulation time and their residence period. The analysis of the MD simulations indicated the simulated WT1 cluster to represent the most stable FGF2/α1-subCD3 interface showing a broad range of interactions together with three ion pairs, E525– K60, E498–R30 and K524–D27. Furthermore, K54 was found to be important as it stabilized the interaction between E498 and R30. As opposed to the WT1, the contact analysis for the WT2 system did not show stable contacts as revealed by MD simulations. In this case, only one pair of residues (K533–M150) was found with a probability of contacts higher than 50%. This suggests that the WT2 system is unlikely to be physiologically relevant. Similarly, the WT3 system appears to be unlikely to represent a relevant interface between FGF2 and α1-subCD3. While it was found to be stable based on MD simulations and its contact map showed contact frequencies of >50%, we did not find K54 and K60 to play a crucial role in stabilizing this potential protein-protein interaction interface. Despite the fact that the WT3 structure in the third cluster was characterized by a relatively large interaction interface, only one ion pair was found to stabilize this interface. In the case of the WT4 system, K54 and K60 were found to play a pivotal role as well as they made up about 25% of the whole contact interface between FGF2 and α1-subCD3. In this case, with E397– K54, E542–K60, E542-K153, and E548– K153, four electrostatic pairs were found to be involved in stabilizing the interaction between FGF2 and α1-subCD3. However, the network of molecular interactions in the WT4 system was not as extended as compared to observations in the WT1 system. In conclusion, the *in silico* results suggest the WT1 structure as the most probable protein-protein interaction surface between FGF2 and α1-subCD3 (Fig. 8).

The atomistic molecular dynamics (MD) simulations were performed using the CHARMM36m ^57^ force field for lipids and proteins, the CHARMM TIP3P force field for water, and the standard CHARMM36 for ions. The GROMACS 2018.3 simulation package ^58^ was used in all simulations. For FGF2, we used its truncated structure [PDB ID: 1BFF; ^59^] from residue 26 to 154 in its monomeric form and the modeled version of human α1-subCD3. The N- and C-terminal groups were modeled as charged residues. The most representative structures of the four clusters were energy-minimized in vacuum using the steepest descent algorithm. The systems were first hydrated and neutralized by an appropriate number of counter-ions, followed by addition of 150 mM potassium chloride to mimic the experimental conditions. All systems were energy-minimized and an equilibration step was used to keep the temperature, pressure and the number of particles constant (NpT ensemble). During this step, proteins were restrained in all dimensions. For the production runs, all atoms in the region involved in the truncation part of α1-subCD3 to the rest of the Na,K-ATPase were restrained in all directions with a force constant of 1000 kJ/mol to avoid the unfolding of the α1-subCD3 domain. A second layer of restraints was applied to the alpha carbons of the residues in the FGF2/α1-subCD3 interaction interface and the thoroughly restrained part. No restraints were employed in the residues involved in the FGF2/α1-subCD3 interaction region. The Nose-Hoover thermostat ^60^ was used to maintain the temperature at 310 K with a time constant of 1.0 ps. The pressure of 1 atm was kept constant using the Parrinello-Rahman barostat ^61^ with a time constant set to 5.0 ps and isothermal compressibility to a value of 4.5 × 10^-5^ bar^-1^. The isotropic pressure-coupling scheme was used. For neighbor searching, we used the Verlet scheme with an update frequency of once every 20 steps. Electrostatic interactions were calculated using the Particle Mesh Ewald method ^62^. Periodic boundary conditions were applied in all directions. The simulations were carried out using an integration time step of 2 fs until they reached 200 ns (Table 3). All analyses were done for the last 100 ns of the 200 ns long simulation trajectories (unless stated otherwise) using standard GROMACS tools and in-house scripts. Finally, using the wild-type structures (WT1-WT4) as a basis, we constructed mutated structures for the mutation of K54 and K60 to E (systems M1-M4) by using the CHARMM-GUI web server.

**Table 3.** Outline of the systems explored through atomistic MD simulations. The wild-type systems WT1-WT4 are the most representative structures of clusters 1-4, respectively, listed in Table 2. The mutated systems M1-M4 were prepared in respective order from WT1-WT4 using CHARMM-GUI. Three replicates were simulated for each of the 8 systems.

| *System* | *# FGF2* | *# SubCD3* | *# Water* | *# K+* | *# Cl-* | *Time (ns)* |
| --- | --- | --- | --- | --- | --- | --- |
| WT1 | 1 | 1 | 42922 | 116 | 118 | 200 |
| WT2 | 1 | 1 | 42922 | 116 | 118 | 200 |
| WT3 | 1 | 1 | 42922 | 116 | 118 | 200 |
| WT4 | 1 | 1 | 42922 | 116 | 118 | 200 |
| M1 | 1 | 1 | 42922 | 118 | 116 | 200 |
| M2 | 1 | 1 | 42922 | 118 | 116 | 200 |
| M3 | 1 | 1 | 42922 | 118 | 116 | 200 |
| M4 | 1 | 1 | 42922 | 118 | 116 | 200 |

### Single molecule TIRF microscopy to study FGF2 membrane recruitment and translocation to the cell surface

Widefield fluorescence and TIRF images were acquired using an Olympus IX81 xCellence TIRF microscope equipped with an Olympus PLAPO 100x/1.45 Oil DIC objective lens and a Hamamatsu ImagEM Enhanced (C9100-13) camera. GFP fluorescence was excited with an Olympus 488 nm, 100 mW diode laser. Data were recorded and exported in Tagged Image File Format (TIFF) and analyzed via Fiji ^63^.

For the quantification of FGF2-GFP recruitment at the inner leaflet of the plasma membrane, cells were seeded in µ-Slide 8 Well Glass Bottom (ibidi) 24 h before live cell imaging experiments. The quantification of FGF2-GFP particles recruitment to the plasma membrane was achieved through the analysis of time-lapse TIRF videos. The frame of each cell was selected by widefield imaging. The number of FGF2-GFP particles was normalized to the cell surface area (μm^2^) and to the expression level of FGF2-GFP. The relative expression levels were quantified for each analyzed cell by measuring the mean intensity of GFP fluorescence at the first frame of each image sequence using ImageJ. The total number of FGF2-GFP particles per cell was quantified using the Fiji plugin TrackMate ^64^. Background fluorescence was subtracted in all the representative images and videos shown.

### Quantification of FGF2-GFP secretion from cells using cell surface biotinylation

Quantification of FGF2-GFP on cell surfaces was done as described previously ^20, 23, 43^. Stable CHO K1 cell lines expressing various forms of FGF2-GFP in a doxycycline-dependent manner were cultured in α-MEM medium supplemented with 10% FCS, 2 mM glutamine, 100 U/ml penicillin and 100 μg/ml streptomycin at 37°C in the presence of 5% CO_2_. Cells were seeded at 0.8 x 10^5^ cells/ml in 6-well plates (Corning Costar). Following 24 hours of incubation, 1 μg/ml of doxycycline (Clontech) was added to induce FGF2-GFP expression. After a further 18 hours of incubation, cells were washed twice with PBS supplemented with 1 mM MgCl_2_ and 0.1 mM CaCl_2_ and incubated with 1 mg/ml of a membrane-impermeable biotinylation reagent (EZ-Link Sulfo-NHS-SS-Biotin, Pierce; dissolved in 150 mM NaCl, 10 mM triethanolamine, pH 9.0, 2 mM CaCl_2_) for 30 min at 4°C. Following one washing step and 20 min of incubation with 100 mM glycine (dissolved in PBS supplemented with 1 mM MgCl_2_ and 0.1 mM CaCl_2_) at 4°C, cells were washed twice with PBS and lysed at 37°C with 1% Nonidet P-40 [in 50 mM Tris/HCl, pH 7.5, 62.5 mM EDTA pH 8, 0.4% deoxycholate, protease inhibitor mixture (Roche Applied Science)]. Insoluble material was removed by centrifugation (10 min, 18000 x g, 4°C). Aliquots from the total cell lysate were taken and the remaining cell lysate was incubated with streptavidin beads (UltraLink immobilized streptavidin; Pierce) for one hour at room temperature. Bound material was eluted with SDS sample buffer for 10 min at 95°C. Total cell lysate and the biotinylated fraction were analyzed by Western blotting using affinity-purified anti-GFP antibodies and monoclonal anti-GAPDH antibodies (Lifetech-Ambion) as primary antibodies and fluorophore-labeled secondary antibodies. Antigen signals were quantified using the Odyssey^®^ CLx Imaging System (LI-COR Biosciences).

**Fig. 4 – supplement 1.**
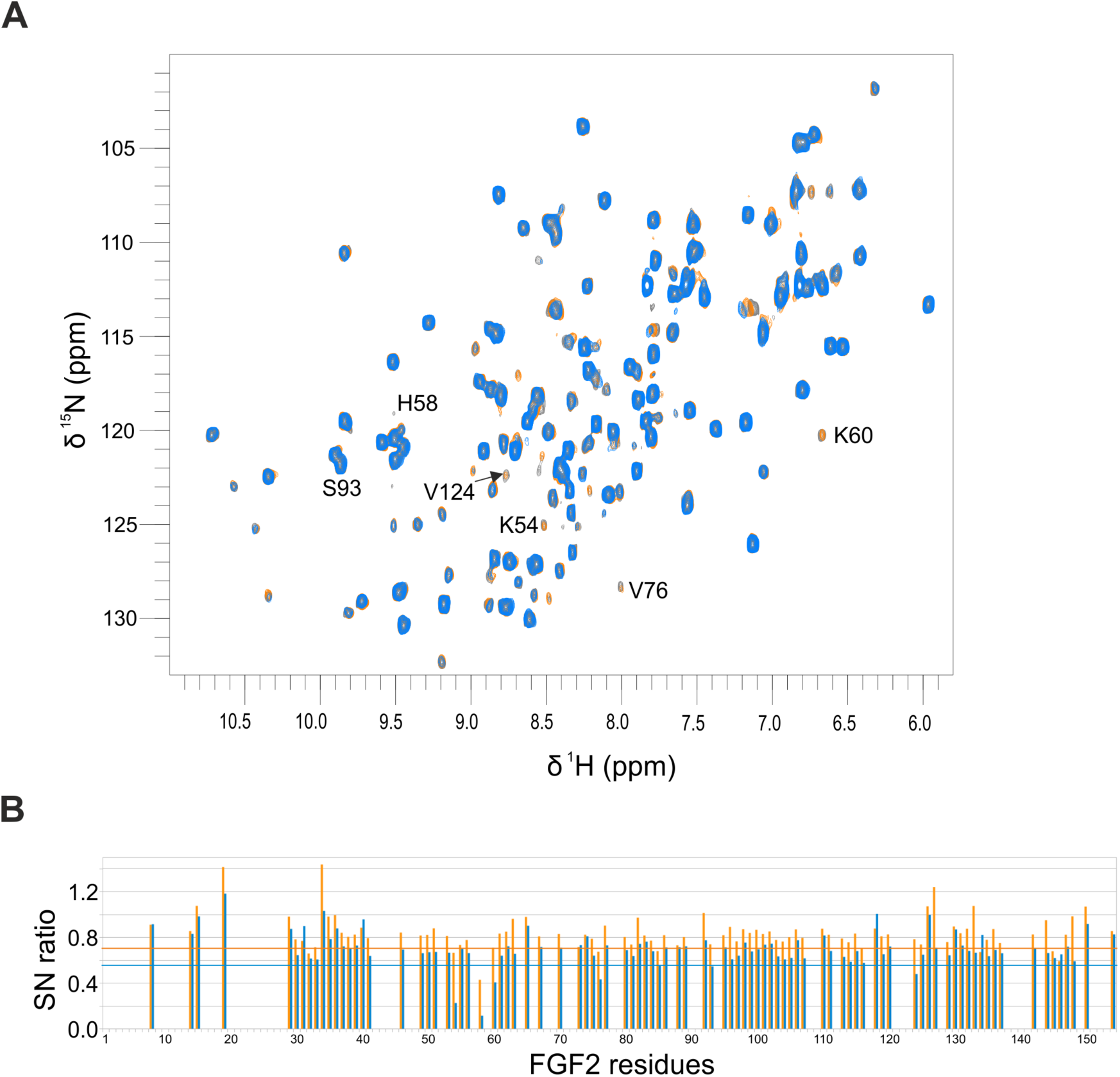
A) Shown is the overlay of the ^1^H-^15^N-HSQC spectra of 77 µM ^15^N-labeled FGF2-C77/95S in the absence (grey) or presence of 70 µM (orange) or 138 µM α1-subCD3 (blue). Chemical shift differences of assigned peaks are very small (<0.04ppm), with an average value of 0.007 ppm. Peak assignments are indicated for those peaks showing a reduction in signal intensity upon addition of 138 µM α1-subCD3 that is considered significant (see also B). B) Plotted are the signal-to-noise ratios (SN ratio) for all assigned peaks with SN(+70 µM α1-subCD3)/SN(-α1-subCD3) in orange and SN(+138 µM α1-subCD3)/SN(-α1-subCD3) in blue. Addition of α1-subCD3 leads to an overall reduction in SN to on average 0.84 (+70 µM) and to 0.70 (+138 µM). A reduction in signal-to-noise beyond the average value minus standard deviation is considered significant. This value is 0.70 for the addition of 70 µM α1 (orange line) and 0.55 for the addition of 138 µM α1 (blue line). Overlapping peaks were omitted from the analysis.

**Fig. 7 – supplement 1:**
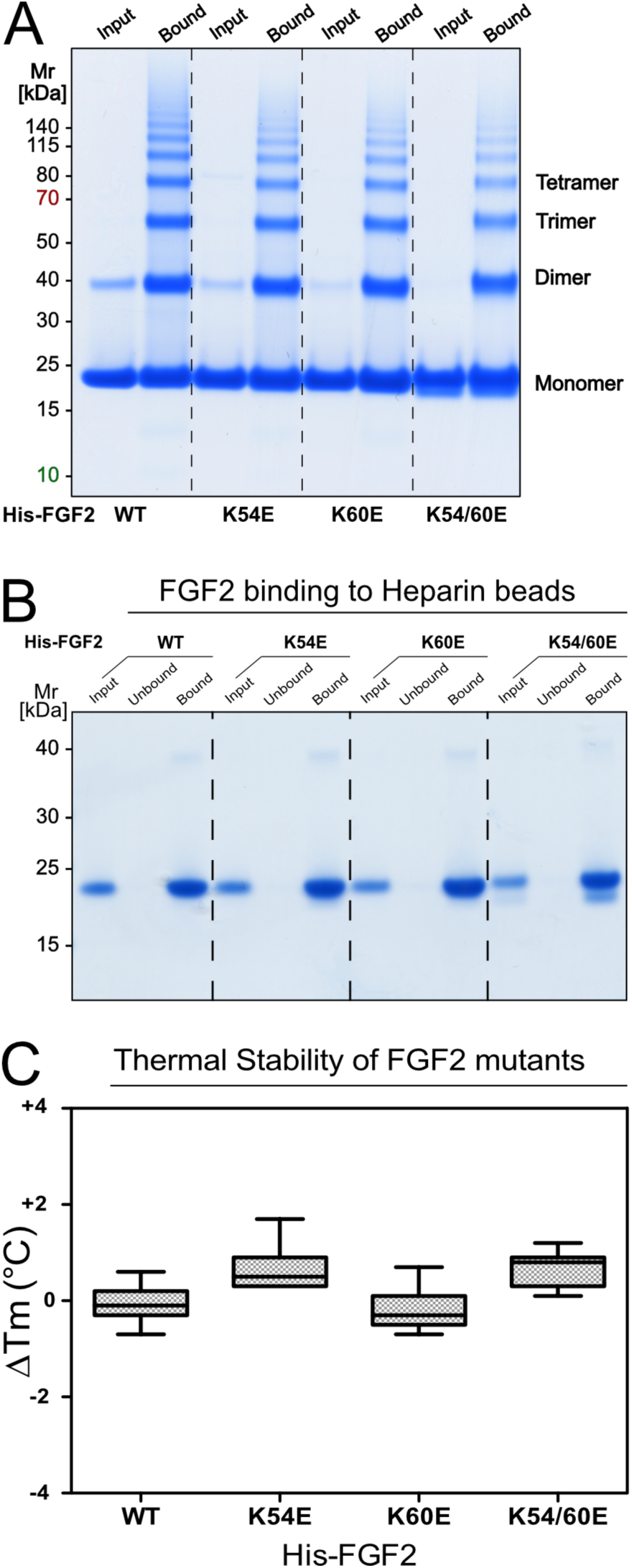
A) Analysis of PI(4,5)P_2_-dependent oligomerization of the FGF2 variant forms indicated. For each FGF2 form, both input material and material bound to PI(4,5)P_2_-containing liposomes is shown. Samples were analyzed by non-reducing SDS-PAGE and Coomassie InstantBlue protein staining. For details, see Materials and Methods. B) Analysis of FGF2 binding to heparin for the variant forms indicated. Following incubation with heparin beads, samples were washed extensively and protein bound to beads was analyzed by reducing SDS-PAGE and Coomassie InstantBlue protein staining. Input (2.5%), Unbound material (2.5%) and bound material (33%) is shown for each of the FGF2 variant forms indicated. For details, see Materials and Methods. C) Analysis of protein folding measuring thermal stability of FGF2 variant forms. Protein samples of 10 µl at a final concentration of 3 mg/ml were analyzed by differential scanning fluorimetry (nanoDSF; ^47^). Deviations from the melting temperature (ΔTm) of FGF2-wt (∼46°C) for each of the FGF2 variant forms indicated were determined with two technical and four biological replicates. Standard deviations are shown. For details, see Materials and Methods.

*Video S1: Atomistic molecular dynamics simulations of the FGF2/α1-subCD3 interface based on the WT1 system.*

Comparison of the interaction of FGF2-wt and FGF2-K54/60E with α1-subCD3 using MD simulations. While the FGF2-wt/α1-subCD3 interface (left panel) remains stable during the whole simulation time (only the last 100 ns are shown in the video), FGF2-K54/60E dissociates from the α1-subCD3 domain already during the first 100 ns of simulation. See Materials and Methods for simulation details. The color coding and the representation styles for FGF2 and α1-subCD3 correspond to Figure 8. In addition, the licorice style is used to highlight the residues that are in contact with K54 and K60 from FGF2.

*Video S2: Single particle analysis of FGF2 membrane recruitment at the inner plasma membrane leaflet in living cells*

Time-lapse TIRF videos (100 ms/frame) of CHO-K1 cell lines expressing the FGF2-GFP fusion proteins introduced in Figs. 10A and 11A (FGF2-K54E, FGF-K60E and FGF2-K54/60E). Background fluorescence was subtracted. Scale bar = 6 µm.

*Video S3: Single particle analysis of FGF2 membrane recruitment at the inner plasma membrane leaflet in living cells*

Time-lapse TIRF videos (100 ms/frame) of CHO-K1 cell lines expressing the FGF2-GFP fusion proteins introduced in Figs. 10B and 11B (FGF2-K54E, FGF-K60E and FGF2-K54/60E combined with K127Q/R128Q/K133Q). Background fluorescence was subtracted. Scale bar = 6 µm.

## References

1. Palade, G. Intracellular aspects of the process of protein synthesis. Science 189, 347–58 (1975).

2. Rothman, J.E. Mechanisms of intracellular protein transport. Nature 372, 55–63 (1994).

3. Rothman, J.E. & Wieland, F.T. Protein sorting by transport vesicles. Science 272, 227–34 (1996).

4. Schekman, R. & Orci, L. Coat proteins and vesicle budding. Science 271, 1526–33 (1996).

5. Dimou, E. & Nickel, W. Unconventional mechanisms of eukaryotic protein secretion. Curr Biol 28, R406–R410 (2018).

6. Rabouille, C. Pathways of Unconventional Protein Secretion. Trends Cell Biol 27, 230–240 (2017).

7. Steringer, J.P. & Nickel, W. A direct gateway into the extracellular space: Unconventional secretion of FGF2 through self-sustained plasma membrane pores. Semin Cell Dev Biol (2018).

8. Akl, M.R. et al. Molecular and clinical significance of fibroblast growth factor 2 (FGF2 /bFGF) in malignancies of solid and hematological cancers for personalized therapies. Oncotarget 7, 44735–44762 (2016).

9. Beenken, A. & Mohammadi, M. The FGF family: biology, pathophysiology and therapy. Nat Rev Drug Discov 8, 235–53 (2009).

10. La Venuta, G., Zeitler, M., Steringer, J.P., Müller, H.M. & Nickel, W. The Startling Properties of Fibroblast Growth Factor 2: How to Exit Mammalian Cells without a Signal Peptide at Hand. J Biol Chem 290, 27015–20 (2015).

11. Brough, D., Pelegrin, P. & Nickel, W. An emerging case for membrane pore formation as a common mechanism for the unconventional secretion of FGF2 and IL-1beta. J Cell Sci 130, 3197–3202 (2017).

12. Dimou, E. et al. Single event visualization of unconventional secretion of FGF2. J Cell Biol (2019).

13. Schäfer, T. et al. Unconventional secretion of fibroblast growth factor 2 is mediated by direct translocation across the plasma membrane of mammalian cells. J Biol Chem 279, 6244–51 (2004).

14. Nickel, W. Unconventional secretory routes: direct protein export across the plasma membrane of mammalian cells. Traffic 6, 607–14 (2005).

15. Zacherl, S. et al. A direct role for ATP1A1 in unconventional secretion of fibroblast growth factor 2. J Biol Chem 290, 3654–65 (2015).

16. Ebert, A.D. et al. Tec-kinase-mediated phosphorylation of fibroblast growth factor 2 is essential for unconventional secretion. Traffic 11, 813–26 (2010).

17. Steringer, J.P. et al. Phosphatidylinositol 4,5-Bisphosphate (PI(4,5)P2)-dependent Oligomerization of Fibroblast Growth Factor 2 (FGF2) Triggers the Formation of a Lipidic Membrane Pore Implicated in Unconventional Secretion. J Biol Chem 287, 27659–69 (2012).

18. Temmerman, K. et al. A direct role for phosphatidylinositol-4,5-bisphosphate in unconventional secretion of fibroblast growth factor 2. Traffic 9, 1204–17 (2008).

19. Temmerman, K. & Nickel, W. A novel flow cytometric assay to quantify interactions between proteins and membrane lipids. J Lipid Res (2009).

20. Zehe, C., Engling, A., Wegehingel, S., Schafer, T. & Nickel, W. Cell-surface heparan sulfate proteoglycans are essential components of the unconventional export machinery of FGF-2. Proc Natl Acad Sci U S A 103, 15479–84 (2006).

21. Nickel, W. Unconventional secretion: an extracellular trap for export of fibroblast growth factor 2. J Cell Sci 120, 2295–9 (2007).

22. Steringer, J.P. et al. Key steps in unconventional secretion of fibroblast growth factor 2 reconstituted with purified components. Elife 6, e28985 (2017).

23. Müller, H.M. et al. Formation of Disulfide Bridges Drives Oligomerization, Membrane Pore Formation and Translocation of Fibroblast Growth Factor 2 to Cell Surfaces. J Biol Chem 290, 8925–8937 (2015).

24. Backhaus, R. et al. Unconventional protein secretion: membrane translocation of FGF-2 does not require protein unfolding. J Cell Sci 117, 1727–36 (2004).

25. Torrado, L.C. et al. An intrinsic quality-control mechanism ensures unconventional secretion of fibroblast growth factor 2 in a folded conformation. J Cell Sci 122, 3322–9 (2009).

26. Evavold, C.L. et al. The Pore-Forming Protein Gasdermin D Regulates Interleukin-1 Secretion from Living Macrophages. Immunity (2017).

27. Katsinelos, T. et al. Unconventional Secretion Mediates the Trans-cellular Spreading of Tau. Cell Rep 23, 2039–2055 (2018).

28. Martin-Sanchez, F. et al. Inflammasome-dependent IL-1beta release depends upon membrane permeabilisation. Cell Death Differ 23, 1219–31 (2016).

29. Monteleone, M. et al. Interleukin-1beta Maturation Triggers Its Relocation to the Plasma Membrane for Gasdermin-D-Dependent and-Independent Secretion. Cell Rep 24, 1425–1433 (2018).

30. Rayne, F. et al. Phosphatidylinositol-(4,5)-bisphosphate enables efficient secretion of HIV-1 Tat by infected T-cells. EMBO J 29, 1348–62 (2010).

31. Zeitler, M., Steringer, J.P., Muller, H.M., Mayer, M.P. & Nickel, W. HIV-Tat Protein Forms Phosphoinositide-dependent Membrane Pores Implicated in Unconventional Protein Secretion. J Biol Chem 290, 21976–84 (2015).

32. Merezhko, M. et al. Secretion of Tau via an Unconventional Non-vesicular Mechanism. Cell Rep 25, 2027–2035 e4 (2018).

33. Florkiewicz, R.Z., Anchin, J. & Baird, A. The inhibition of fibroblast growth factor-2 export by cardenolides implies a novel function for the catalytic subunit of Na+,K+-ATPase. J Biol Chem 273, 544–51 (1998).

34. Kaplan, J.H. Biochemistry of Na,K-ATPase. Annu Rev Biochem 71, 511–35 (2002).

35. Engling, A. et al. Biosynthetic FGF-2 is targeted to non-lipid raft microdomains following translocation to the extracellular surface of CHO cells. J. Cell Sci. 115, 3619–3631 (2002).

36. Dahl, J.P., Binda, A., Canfield, V.A. & Levenson, R. Participation of Na,K-ATPase in FGF-2 Secretion: Rescue of Ouabain-Inhibitable FGF-2 Secretion by Ouabain-Resistant Na,K-ATPase alpha Subunits. Biochemistry 39, 14877–14883 (2000).

37. Agostini, S. et al. Inhibition of Non Canonical HIV-1 Tat Secretion Through the Cellular Na+,K+-ATPase Blocks HIV-1 Infection. EBioMedicine 21, 170–181 (2017).

38. Nickel, W. & Seedorf, M. Unconventional mechanisms of protein transport to the cell surface of eukaryotic cells. Annu Rev Cell Dev Biol 24, 287–308 (2008).

39. Nickel, W. & Rabouille, C. Mechanisms of regulated unconventional protein secretion. Nat Rev Mol Cell Biol 10, 148–55 (2009).

40. Hilge, M. et al. ATP-induced conformational changes of the nucleotide-binding domain of Na,K-ATPase. Nat Struct Biol 10, 468–74 (2003).

41. La Venuta, G. et al. Small Molecule Inhibitors Targeting Tec Kinase Block Unconventional Secretion of Fibroblast Growth Factor 2. J Biol Chem 291, 17787–803 (2016).

42. Moy, F.J., Seddon, A.P., Campbell, E.B., Bohlen, P. & Powers, R. 1H, 15N, 13C and 13CO assignments and secondary structure determination of basic fibroblast growth factor using 3D heteronuclear NMR spectroscopy. J Biomol NMR 6, 245–54 (1995).

43. Seelenmeyer, C. et al. Cell surface counter receptors are essential components of the unconventional export machinery of galectin-1. J Cell Biol 171, 373–81 (2005).

44. Nickel, W. The unconventional secretory machinery of fibroblast growth factor 2. Traffic 12, 799–805 (2011).

45. Rabouille, C., Malhotra, V. & Nickel, W. Diversity in unconventional protein secretion. J Cell Sci 125, 5251–5 (2012).

46. Steringer, J.P., Müller, H.M. & Nickel, W. Unconventional secretion of fibroblast growth factor 2--a novel type of protein translocation across membranes? J Mol Biol 427, 1202–10 (2015).

47. Alexander, C.G. et al. Novel microscale approaches for easy, rapid determination of protein stability in academic and commercial settings. Biochim Biophys Acta 1844, 2241–50 (2014).

48. Vranken, W.F. et al. The CCPN data model for NMR spectroscopy: development of a software pipeline. Proteins 59, 687–96 (2005).

49. Chojnacki, S., Cowley, A., Lee, J., Foix, A. & Lopez, R. Programmatic access to bioinformatics tools from EMBL-EBI update: 2017. Nucleic Acids Res 45, W550–W553 (2017).

50. Morth, J.P. et al. Crystal structure of the sodium-potassium pump. Nature 450, 1043–9 (2007).

51. Lee, J. et al. CHARMM-GUI Input Generator for NAMD, GROMACS, AMBER, OpenMM, and CHARMM/OpenMM Simulations Using the CHARMM36 Additive Force Field. J Chem Theory Comput 12, 405–13 (2016).

52. Chaudhury, S. & Gray, J.J. Conformer selection and induced fit in flexible backbone protein-protein docking using computational and NMR ensembles. J Mol Biol 381, 1068–87 (2008).

53. Gray, J.J. et al. Protein-protein docking with simultaneous optimization of rigid-body displacement and side-chain conformations. J Mol Biol 331, 281–99 (2003).

54. Wang, C., Bradley, P. & Baker, D. Protein-protein docking with backbone flexibility. J Mol Biol 373, 503–19 (2007).

55. Wang, C., Schueler-Furman, O. & Baker, D. Improved side-chain modeling for protein-protein docking. Protein Sci 14, 1328–39 (2005).

56. Daura, X. et al. Peptide folding: Ehen simulation meets experiment. Angew Chem Int Ed Engl 38, 236–240 (1999).

57. Huang, J. et al. CHARMM36m: an improved force field for folded and intrinsically disordered proteins. Nat Methods 14, 71–73 (2017).

58. Abraham, M., et al. GROMACS: High performance molecular simulations through multi-level parallelism from laptops to supercomputers. SoftwareX 1(2015).

59. Kastrup, J.S., Eriksson, E.S., Dalboge, H. & Flodgaard, H. X-ray structure of the 154-amino-acid form of recombinant human basic fibroblast growth factor. comparison with the truncated 146-amino-acid form. Acta Crystallogr D Biol Crystallogr 53, 160–8 (1997).

60. Evans, D.J. & Holian, B.L. The Nose–Hoover thermostat. J. Chem. Phys. 83, 4096 (1985).

61. Parrinello, M. & Rahman, A. Polymorphic transitions in single crystals: A new molecular dynamics method. J. Appl. Phys. 52, 7182–7190 (1981).

62. Darden, T., York, D. & Pedersen, L. Particle mesh Ewald: An N-log(N) method for Ewald sums in large systems. J. Chem. Phys. 98, 10089–10092 (1993).

63. Schindelin, J., et al. Fiji: an open-source platform for biological-image analysis. Nat Methods 9, 676–82 (2012).

64. Tinevez, J.Y. et al. TrackMate: An open and extensible platform for single-particle tracking. Methods 115, 80–90 (2017).

